# *Plasmodium falciparum* population genetic complexity influences transcriptional profile and immune recognition of highly related genotypic clusters

**DOI:** 10.1101/2020.01.03.894220

**Authors:** Amy K. Bei, Daniel B. Larremore, Kazutoyo Miura, Ababacar Diouf, Nicholas K. Baro, Rachel F. Daniels, Allison Griggs, Eli L. Moss, Daniel E. Neafsey, Awa B. Deme, Mouhamad Sy, Stephen Schaffner, Ambroise D. Ahouidi, Daouda Ndiaye, Tandakha Dieye, Souleymane Mboup, Caroline O. Buckee, Sarah K. Volkman, Carole A. Long, Dyann F. Wirth

**Affiliations:** Department of Immunology & Infectious Diseases, Harvard TH Chan School of Public Health, Boston, MA, USA; Laboratory of Bacteriology and Virology, Le Dantec Hospital, Faculty of Medicine and Pharmacy, Cheikh Anta Diop University, Dakar, Senegal; Laboratory of Parasitology and Mycology, Faculty of Medicine and Pharmacy, Cheikh Anta Diop University, Dakar, Senegal; Yale School of Public Health, Laboratory of Epidemiology and Public Health, New Haven, USA; Department of Computer Science, University of Colorado Boulder, Boulder, CO, USA; BioFrontiers Institute, University Colorado at Boulder, Boulder, CO, USA; Department of Epidemiology, Harvard TH Chan School of Public Health, Boston, MA, USA; Center for Communicable Disease Dynamics, Harvard TH Chan School of Public Health, Boston, Massachusetts, USA; Santa Fe Institute, 1399 Hyde Park Road, Santa Fe, NM, USA; Laboratory of Malaria and Vector Research, National Institute of Allergy and Infectious Diseases, National Institutes of Health, Bethesda, MD, USA; Broad Institute of MIT and Harvard, 7 Cambridge Center, Cambridge, MA 02142, USA; Institut de Recherche en Santé, de Surveillance Epidemiologique et de Formations, Dakar, Senegal

## Abstract

As transmission intensity has declined in Senegal, so has the genetic complexity of circulating *Plasmodium falciparum* parasites, resulting in specific genotypes emerging and persisting over years. We address whether changes in parasite genetic signatures can alter the immune repertoire to variant surface antigens, and whether such responses can influence the expansion or contraction of specific parasite genotypes in the population. We characterize parasites within genotypic clusters, defined as identical by a 24-SNP molecular barcode and a haplotype identifier for other highly polymorphic loci; we measure expression of variant surface antigens (VSA) such as PfEMP-1 by transcript expression typing and expressed *var* DBL1*α* sequencing in ex vivo and short-term adapted RNA samples; and we measure IgG responses against VSAs from short-term adapted parasites. We find that parasites within genotypic clusters are genetically identical at other highly polymorphic loci. These parasites express similar Ups *var* classes and largely the same dominant *var* DBL1*α* sequences ex vivo. These parasites are recognized similarly by anti-VSA antibodies after short-term adaptation to culture; however, antibody responses do not correlate with genotype frequencies over time. Both genotype-specific and multiple genotype-reactive surface IgG responses are observed in this population. Parasites with identical genomes are extremely similar in their expression and host antibody recognition of VSAs. Monitoring changes in population-level parasite genomics and transmission dynamics is critical, as fluctuations will influence the breadth of resulting host immune responses to circulating parasite genotypes. These findings suggest shared immune recognition of genetically similar parasites, which has implications for both our understanding of immunity and vaccine development strategies in malaria elimination settings.

## INTRODUCTION

Great efforts have been made in recent years to control, eliminate, and eventually eradicate malaria, a devastating tropical disease caused by parasites of the genus Plasmodium. Despite great progress, malaria is still a public health priority and remains one of the leading causes of death in children under the age of 5 in sub-Saharan Africa [1]. As successful malaria control measures have been implemented, transmission intensity in some regions has declined [1, 2]. In Senegal, this drop in transmission intensity has been accompanied by corresponding decreases in the multiplicity of infection and genotypic diversity in the Plasmodium falciparum population. This decreased diversity is characterized by the emergence of clusters of parasites that share a common genetic signature (CGS) [3]. CGS clusters are defined by a genetic barcode consisting of 24 Single Nucleotide Polymorphisms (SNPs) [4, 5], and have been tracked along with overall transmission intensity in Thiès, Senegal for the past thirteen years. Initially, declines in transmission correlated with decreased parasite genotypic richness [4]. More recently, however, allelic variation has rebounded [3], raising questions about the complex relationships between the dynamics of transmission, the genetic diversity of the parasite population, and the human immune responses at individual and population scales.

Individuals who survive a malaria infection develop a broadly protective but non-sterilizing immunity. This is due to the parasite’s evolved mechanism for escape from immune clearance, in part through antigenic variation of proteins exposed on the surface of infected erythrocytes [6, 7], which are collectively referred to as variant surface antigens (VSA). The level of exposure to specific antigen variants influences the degree of immune protection [8–11], so as exposure to the diverse repertoire of parasite antigens decreases, the corresponding immune responses also become more restricted in their recognition. This, in turn, gives rise to the possibility of immune escape and fluctuations in both population-level parasite genotypic richness and the degree of immune protection. However, it is unclear whether this possible mechanism explains the patterns observed in Senegal, and whether those patterns are likely to generalize to other low-transmission and pre-elimination settings.

#### Plain Text Summary

A key virulence determinant of Plasmodium falciparum is its capacity for genetic diversity allowing parasites to develop resistance to antimalarial drugs and rendering the parasite “invisible” to natural immunity, due to variation in the antigens recognized by the immune system. As transmission intensity declines, population-level genetic complexity decreases as well, resulting in the emergence of highly-related genotypic clusters. In Senegal, we studied multiple highly-related parasite clusters and find that gene expression and antibody recognition of variant surface antigens are highly similar within clusters: the more related the parasites are genome-wide, the more similar the immune response. As malaria control priorities shift toward elimination, interactions between genetic diversity and immunological recognition are critical to monitor in regions with changing transmission dynamics.

We previously studied the relationships between CGS clusters, their antigens, and population immunity across the two years prior to the diversity rebound. We found that immune responses to the VSAs of parasites within a single CGS cluster were statistically more similar than controls, and further found that those parasites also express statistically more similar *var* genes, which encode a dominant VSA PfEMP-1 [12]. Furthermore, these same CGS parasites dropped from high to low prevalence over the two-year study, while immune recognition of their VSAs correspondingly rose over the same period. Taken together, these patterns suggest that CGS clusters with highly similar VSA profiles are under frequencydependent selection by human population-level immunity. However, with data from only one genotypic cluster studied over two years, collected prior to the diversity rebound, more fundamental rules governing the relationship between changes in parasite population genotypic heterogeneity and host population immune responses have yet to be fully uncovered.

In this paper, we advance our understanding of these relationships by reporting an extended study of seven different clusters of CGS parasites from Thiès, spanning seven transmission seasons from 2007 to 2014. In particular, we analyze VSAs and human immunity over time, characterizing parasites from the barcode-defined CGS clusters by sequencing other polymorphic loci and genomic *var* repertoires, and further analyzing *var* transcript expression and immune recognition by VSA flow cytometry. Because the CGS clusters we study have different dynamics of emergence, growth, and decline, we explore how changes in genotypic richness can alter human immune repertoires to variant surface antigens, and how such altered immune repertoires could, in turn, influence or modulate the expansion or contraction of specific parasite genotypes. Thus, by combining genetic, transcriptional, immunological, and host-pathogen interaction data, this study represents the most comprehensive analysis of parasites persistence dynamics to date. Such findings have important implications for how fluctuations in population-level parasite genetic diversity can affect specific immune responses and are critical to our understanding of both naturally acquired protective immune responses as well as vaccine-induced immune responses to specific parasite variants.

## RESULTS

### Genotypic clusters of CGS parasites show diverse patterns of emergence, persistence, and decline, but no spatial or clinical correlations

During the seven transmission seasons under study, parasite population structure in Thiès, Senegal was determined by applying an established 24-SNP molecular barcode to samples from single-clone infections collected by passive case detection [3]. At the level of this barcode, parasites cluster definitively into numerous haplotypes which we call Common Genetic Signature (CGS) groups or clusters [3]. Plotting the prevalence of CGS Haplotypes between 2007 and 2014 reveals a diverse set of emergence, persistence, and decline dynamics (Figure 1, Table S1). For instance, Haplotype 29 emerged in 2007 and persisted at high overall frequencies until 2012, with continued persistence through 2014. In contrast, Haplotypes 21, 24, 26, and 99, emerged and then disappeared rapidly, spanning just two transmission seasons. Still others, such as Haplotype 81, emerged and occured in just a single year and did not persist beyond the single transmission season, while Haplotype 728 emerged at the end of the study period. Finally, some parasites did not cluster with others, which we designate as “unique,” and include in our study to provide a contrasting control for parasites within clusters (Figure 1B). We selected CGS clusters for further in-depth study, motivated by two broad questions. First, how similar are parasites from the same CGS cluster to each other, at both the genetic and transcription level? Second, how do Haplotype persistence and frequency influence the immune response to such clusters in the population? Based on these questions, we selected a representative sample of the diversity of emergence, persistence, and decline dynamics consisting of Haplotypes 24, 26, 29, 728, and a set of haplotypes that were only observed once, which are hereafter referred to as “unique.” Spatio-temporal locations of these CGS Haplotypes were mapped, revealing no obvious clustering patterns by genotype (Figure S1). There were no specific clinical parameters such as patient age, temperature, parasitemia, etc. that differed significantly by cluster.

**FIG. 1.**
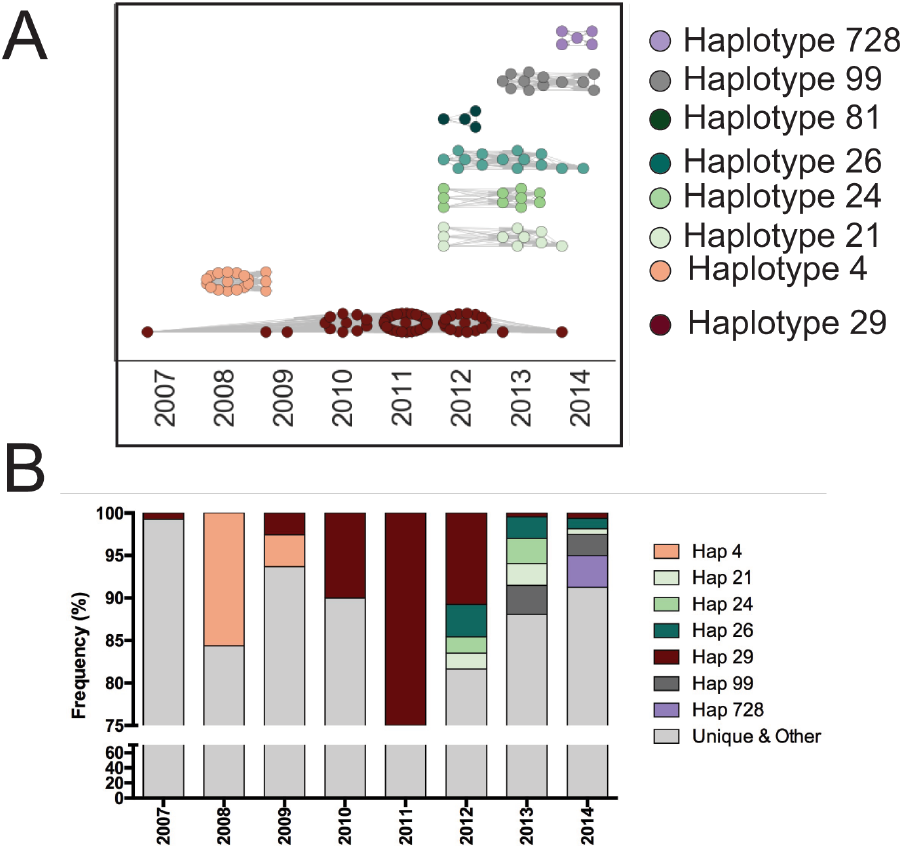
Patterns of parasite persistence over time. A. Parasite genotypes (haplotypes) based on the 100% similarity of the molecular barcode from 2007 to 2014. Each dot represents an individual parasite isolate. Haplotypes investigated in depth in this study, or in past studies discussed in this study, are highlighted to the right.B. The specific frequencies of each CGS Haplotype are shown as the percentage of all sampled single-clone infections in that given year. Grey represents “Unique and Other”: combined Unique haplotypes (haplotypes observed only once from 2007 to present) as well as haplotypes not discussed in this study.

### Barcode-identical CGS parasites are also identical at highly polymorphic loci: CSP, TRAP, and SERA2

In principle, parasites in barcode-identical CGS clusters need not be identical at other loci, since the barcode is based on a set of stable and neutral SNPs, while other, non-neutral loci are highly polymorphic. To this end, we sequenced circumsporozoite protein (CSP), thrombospondin-related anonymous protein (TRAP), and serine repeat antigen 2 (SERA2)—three loci known to be under immune selection—for CGS parasites and found that all parasites within a barcode-identical cluster are also identical at these three polymorphic loci (Figure S2). Furthermore, each distinct cluster contains corresponding distinct haplotypes, with one exception: parasites from Haplotypes 728 and 29 share the same sequence at the TRAP and SERA2 loci but were distinct at the CSP C-terminus. When CSP, TRAP, and SERA2 sequences are combined to form a single haplotype, parasites within clusters are identical, and each cluster contains its own individual composite haplotype (Figure S2D). Furthermore, the parasites designated as unique, which do not belong to any CGS cluster, were all found to have unique CSP, TRAP, and SERA2 haplotypes, and all observed parasites were distinct from the 3D7 reference at these loci. In summary, while there is overall population diversity at these highly polymorphic loci, parasites within CGS clusters are nevertheless identical at CSP, TRAP, and SERA2.

### CGS parasites express similar *var* DBL1*α* sequences and *var* Ups classes ex vivo

We previously observed that CGS parasites from Haplotype 4 express similar *var* genes, which encode PfEMP1, a major variant surface antigen, more than would be expected by chance [3, 12]. We therefore sought to determine if this transcriptional similarity was generalizable to other clusters ex vivo, particularly after finding identical CSP, TRAP, and SERA2 loci. *var* transcriptional profiles are of particular interest due to their association with severe disease phenotypes and antigenic variation. To that end, we analyzed three to four randomly selected parasites from each cluster of interest, focusing on a single year for each, provided that there were more than three independent ex vivo samples (2012: haplotype 24; 2013: haplotypes 26, 29, 99; 2014: haplotype 728, and unique). We evaluated *var* transcription similarity between parasites by analyzing both the sequence similarity of transcribed DBL1*α* domains, and the dominant upstream promoter sequence (Ups) classes (see Methods).

DBL1*α* has been implicated in the phenotypes of rosetting and endothelial binding [15–17], and is present in the majority of *var* genes, in contrast to other domains [18]. At the expressed *var* sequence level, DBL1*α* sequencing of CGS clusters revealed that many, but not all members of a CGS haplotype expressed the same dominant *var* gene (Table I). In addition to the most dominant *var* within clusters, an analysis of overlap between expressed *var* repertoires shows significant overlap among sequences within a cluster (Figure 2). Further, as expected, unique parasites all expressed unique dominant *var* genes and showed no sequence overlap with parasites in CGS clusters, reinforcing the notion that these parasites serve as a control and contrast for parasites within clusters (Figure 2A, Table I; see parasites Uni1 12, Uni4 14). These results hold without modification when Bayesian methods are used to account for uncertainty in repertoire overlap networks, caused by subsampling (Figure S4) [19].

**TABLE I.** **Dominant DBL1***α* **var sequences for each strain from ex vivo RNA**. Expressed *var* DBL1*α* sequences (from cDNA) were compared for parasite strains within CGS clusters to determine whether strains within a cluster have any overlap in the dominant *var* they express ex vivo. Dominant *vars* are listed here according to strain and grouped within cluster (shaded and unshaded). Additionally, *var* identity based on homology both by PSI-blast and the 3D7 genome from PlasmoDB.org are also listed. *Vars* that are similar within clusters are bolded. ^*^ indicate samples with very few high quality sequences, and thus less confidence in the proportions.

| Sample ID code | Haplotype | Sequence Number | Proportion amplified | PSI Blast Genbank Accession | cysteine/ PoLV group | Annotation | PlasmoDB 3D7 |
| --- | --- | --- | --- | --- | --- | --- | --- |
| 29.9 _12 | 29 | B6 | 8/17 (47%) | **CAJ40551.1** | 4 | Kilifi, sequence group 4 | **PF3D7_0421100** |
| 29.9 _12 | 29 | C3 | 5/17 (29%) | ADX61958.1 | 4 | Pikine 38 | **PF3D7_1240400** |
| 29.12 _12 | 29 | A11 | 11/36 (30.6%) | ADX60696.1 | 4 | Kilifi 09 | PF3D7_1255200 |
| 29.12 _12 | 29 | A12 | 8/36 (22.2%) | ADX62021.1 | 4 | Pikine 49 | PF3D7_0426000 |
| 29.12 _12 | 29 | A6 | 7/36 (19.4%) | **CAJ40551.1** | 4 | Kilifi, sequence group 4 | **PF3D7_0421100** |
| 29.12 _12 | 29 | E1 | 14/36 (38.9%) | CAJ39112.1 | 3 | Kilifi, sequence group 3 | PF3D7_0800300 |
| 29.13 _12 | 29 | E11 | 3/6 (50%) | ADX61532.1 | 4 | Pikine 05 | **PF3D7_1240400** |
| 29.14 _12 | 29 | A12 | 18/20 (90%) | ADX61518.1 | 5 | Pikine 04 | **PF3D7_0808700** |
| 728.4 _14 | 728 | A6 | 6/16 (37.5%) | CCC18641.1 |  | Kenya 8344 | PF3D7_0800300 |
| 24.2 _13 | 24 | B6 | 4/31 (12.9%) | ABC95905.1 | 1 | Mali variant HYP5b | PF3D7_1150400 |
| 24.2 _13 | 24 | A2 | 3/31 (9.7%) | CAJ39011.1 | 4 | Kilifi, sequence group 4 | PF3D7_0426000 |
| 24.2 _13 | 24 | A4 | 3/31 (9.7%) | **ABD92278.1** | **2** | East Africa upsAvar | **PF3D7_0800200** |
| 24.2 _13 | 24 | A1 | 3/31 (9.7%) | AAF36641.1 | 4 | Kenya 13 | PF3D7_0712000 |
| 24.4 _13 | 24 | F1 | 4/20 (20%) | **ABD92278.1** | **2** | East Africa upsAvar | **PF3D7_0800200** |
| 99.2 _13 | 99 | F5 | 4/14 (28.6%) | **AHI95490.1** | **4** | Malawi | **PF3D7_1255200** |
| 99.2 _13 | 99 | D5 | 3/14 (21.4%) | **ACH54072.1** | **4** | PNG | **PF3D7_0400100** |
| 99.3 _13 | 99 | B9 | 5/28 (17.9%) | ABC95902.1 | 2 | Mali variant HYP4a | **PF3D7_0600200** |
| 99.3 _13 | 99 | A10 | 4/28 (14.3%) | ACH53778.1 | 1 | PNG | PF3D7_1300300 |
| 99.3 _13 | 99 | B8 | 4/28 (14.3%) | **AIZ74248.1** | **2** | PNG WOS4 | **PF3D7_0600200** |
| 99.3 _13 | 99 | F10 | 4/28 (14.3%) | **ACH54072.1** | **4** | PNG | **PF3D7_0400100** |
| 99.4 _13 | 99 | A5 | 5/29 (17.2%) | CCF76354.1 | 4 | Kilifi 28H | PF3D7_1240400 |
| 99.4 _13 | 99 | C5 | 5/29 (17.2%) | **AIZ74248.1** | **2** | PNG WOS4 | **PF3D7_0600200** |
| 99.5 _13 | 99 | D11 | 10/36 (27.8%) | **AIZ74248.1** | **2** | PNG WOS4 | **PF3D7_0600200** |
| 99.5 _13 | 99 | B11 | 4/36 (11.1%) | **AHI95490.1** | **4** | Malawi | **PF3D7_1255200** |
| 99.5 _13 | 99 | A7 | 6/36 (16.7%) | ACH53959.1 | 1 | PNG | PF3D7_0937600 |
| 26.2 _12 | 26 | C10 | 4/29 (13.8%) | AHI95077.1 | 2 | Malawi | PF3D7_0600200 |
| 26.2 _12 | 26 | D7 | 3/29 (10.3%) | ABD92258.1 | 1 | South Africa upsAvar | **PF3D7_0400400** |
| 26.3 _13 | 26 | B7 | 2/17 (11.8%) | ADX61577.1 | 5 | Pikine 06 | PF3D7_0808600 |
| 26.3 _13 | 26 | A9 | 2/17 (11.8%) | ADX61116.1 | 4 | Kilifi KIL36 | PF3D7_0412400 |
| 26.3 _13 | 26 | F10 | 2/17 (11.8%) | CAJ39120.1 | 1 | Kilifi sequence group 6 | **PF3D7_0400400** |
| 26.3 _13 | 26 | F7 | 2/17 (11.8%) | ADX61765.1 | 1 | Pikine 22 | PF3D7_1300300 |
| 26.4 _13 | 26 | C10 | 2/4 (50%)* | AIZ73177.1 | 1 | PNG MUG27 | PF3D7_0937600 |
| 26.4 _13 | 26 | B10 | 1/4 (25%)* | ADX62024.1 | 4 | Pikine 49 | PF3D7_1255200 |
| 26.4 _13 | 26 | G11 | 1/4 (25%)* | ETW50692.1 | 2 | Mali PS096 | PF3D7_0800300 |
| Uni1 _12 | Unique_12 | C4 | 2/2 (100%)* | AIZ74255.1 | 4 | PNG WOS5 | PF3D7_0412900 |
| Uni4 _14 | Unique_14 | A9 | 5/38 (13.2%) | KOB63545.1 | 4 | HB3 Pfemp1 | PF3D7_0324900 |
| Uni4 _14 | Unique_14 | C11 | 5/38 (13.2%) | ADX60825.1 | 4 | Kilifi KIL22 | PF3D7_1240300 |

**TABLE II.** **Dominant DBL1***α* **var sequences for each strain from in vitro adapted RNA**. Expressed *var* DBL1*α* sequences (from cDNA) were compared for parasite strains within CGS clusters to determine whether strains within a cluster have any overlap in the dominant *var* they express after in vitro adaptation. Dominant *vars* are listed here according to strain and grouped within cluster (shaded and unshaded). Additionally, *var* identity based on homology both by PSI-blast and the 3D7 genome from PlasmoDB.org are also listed. *Vars* that are similar within clusters are bolded. ^*^ indicate samples with very few high-quality sequences, and thus less confidence in the proportions.

| Sample ID code | Haplotype | Sequence Number | Proportion amplified | PSI Blast Genbank Accession | cysteine/ PoLV group | Annotation | PlasmoDB 3D7 |
| --- | --- | --- | --- | --- | --- | --- | --- |
| 29.1_07 | 29 | A4 | 7/38 (18%) | **ADX61958.1** | 4 | Pikine PIK38 | **PF3D7_1240400** |
| 29.1_07 | 29 | A10 | 6/38 (16%) | **AAR32006.1** | 4 | PNG | PF3D7_1219300 |
| 29.1_07 | 29 | A3 | 5/38 (13%) | AAR32021.1 | 4 | PNG | **PF3D7_1255200** |
| 29.2_09 | 29 | E5 | 5/33 (15%) | ACT21635.1 | 4 | Gabon 130_01 | PF3D7_1041300 |
| 29.2_09 | 29 | E7 | 5/33 (15%) | ADX61714.1 | 5 | Pikine PIK12 | PF3D7_0421300 |
| 29.2_09 | 29 | F4 | 4/33 (12%) | **AAR32006.1** | 4 | PNG | **PF3D7_0809100** |
| 29.2_09 | 29 | F3 | 4/33 (12%) | **AIZ74167.1** | 4 | PNG WOS32 | **PF3D7_0412900** |
| 29.3_11 | 29 | A4 | 6/46 (13%) | ADX61472.1 | 4 | Pikine PIK02 | **PF3D7_1255200** |
| 29.3_11 | 29 | B1 | 6/46 (13%) | **ADX61868.1** | 4 | Pikine PIK23 | **PF3D7_1100100** |
| 29.4_11 | 29 | E1 | 14/44 (32%) | AHI95553.1 | 4 | Malawi | PF3D7_1200100 |
| 29.4_11 | 29 | F1 | 8/44 (18%) | **ETW48207.1** | 4 | Mali PS096 | **PF3D7_0413100** |
| 29.4_11 | 29 | E8 | 4/44 (9%) | ASY08685.1 | 2 | Malawi | PF3D7_0600200 |
| 29.6_11 | 29 | A1 | 5/40 (13%) | **ADX61868.1** | 4 | Pikine PIK23 | **PF3D7_1100100** |
| 29.9_12 | 29 | E7 | 9/33 (27%) | ETW36135.1 | 4 | Tanzania | **PF3D7_0809100** |
| 29.9_12 | 29 | E4 | 5/33 (15%) | **AIZ74167.1** | 4 | PNG WOS32 | **PF3D7_0412900** |
| 29.12_12 | 29 | A9 | 6/33 (18%) | **ADX61958.1** | 4 | Pikine PIK38 | **PF3D7_1240400** |
| 29.12_12 | 29 | A3 | 6/33 (18%) | **AIZ74167.1** | 4 | PNG WOS32 | **PF3D7_0412900** |
| 29.12_12 | 29 | A4 | 4/33 (12%) | **ETW48207.1** | 4 | Mali PS096 | **PF3D7_0413100** |
| 29.15_12 | 29 | E7 | 6/41 (15%) | **ADX61958.1** | 4 | Pikine PIK38 | **PF3D7_1240400** |
| 29.15_12 | 29 | F4 | 5/41 (12%) | **ADX61868.1** | 4 | Pikine PIK23 | **PF3D7_1100100** |
| 29.15_12 | 29 | E8 | 5/41 (12%) | **AAR32006.1** | 4 | PNG | **PF3D7_0809100** |
| 29.16_13 | 29 | A1 | 12/40 (30%) | **ADX61868.1** | 4 | Pikine PIK23 | **PF3D7_1100100** |
| 29.16_13 | 29 | A4 | 10/40 (25%) | **AAR32006.1** | 4 | PNG | **PF3D7_0809100** |
| 29.16_13 | 29 | A12 | 6/40 (15%) | **ADX61958.1** | 4 | Pikine PIK38 | **PF3D7_1240400** |
| 24.1_12 | 24 | A6 | 14/40 (35%) | ABC95905.1 | 1 | Mali | PF3D7_1150400 |
| 24.2_13 | 24 | E4 | 8/42 (19%) | CCI79415.1 | 4 | Kenya 9376 | PF3D7_0412400 |
| 24.2_13 | 24 | E3 | 7/42 (17%) | CCF76281.1 | 3 | Kilifi 44D | PF3D7_0800300 |
| 24.2_13 | 24 | E9 | 6/42 (14%) | ADX61323.1 | 4 | Kilifi KIL44 | PF3D7_0400100 |
| 24.2_13 | 24 | E11 | 6/42 (14%) | **AAF36641.1** | 4 | Kenya 13 | **PF3D7_0712000** |
| 24.3_13 | 24 | A9 | 7/43 (16%) | KNC36886.1 | 3 | RAJ116 | PF3D7_0115700 |
| 24.3_13 | 24 | A1 | 7/43 (16%) | AAF36633.1 | 4 | Kenya 16 | PF3D7_0421100 |
| 24.4_13 | 24 | E2 | 4/38 (11%) | CAD60899.1 | 2 | Venezuela VZ2 | PF3D7_0800200 |
| 24.4_13 | 24 | F4 | 4/38 (11%) | AHI95009.1 | 4 | Malawi | PF3D7_0100100 |
| 24.4_13 | 24 | E5 | 4/38 (11%) | **AAF36641.1** | 4 | Kenya 13 | **PF3D7_0712000** |
| 26.1_12 | 26 | A3 | 9/40 (23%) | AHI95077.1 | 2 | Malawi | PF3D7_0600200 |
| 26.1_12 | 26 | A8 | 7/40 (18%) | ADX61367.1 | 4 | Kilifi KIL45 | PF3D7_0412700 |
| 26.3_13 | 26 | E10 | 7/43 (16%) | **ADX61577.1** | 5 | Pikine PIK06 | **PF3D7_0808600** |
| 26.3_13 | 26 | E5 | 5/43 (12%) | **ADX61116.1** | 4 | Kilifi KIL36 | **PF3D7_0412400** |
| 26.3_13 | 26 | E2 | 5/43 (12%) | AIZ73177.1 | 1 | PNG MUG27 | PF3D7_0937600 |
| 26.4_13 | 26 | A3 | 5/41 (12%) | **ADX61116.1** | 4 | Kilifi KIL36 | **PF3D7_0412400** |
| 26.4_13 | 26 | B4 | 5/41 (12%) | **ADX61577.1** | 5 | Pikine PIK06 | **PF3D7_0808600** |
| 26.4_13 | 26 | B1 | 5/41 (12%) | ANJ21274.1 | 1 | Tanzania 1983-15 | PF3D7_0400400 |
| 26.6_13 | 26 | E4 | 16/43 (37%) | **ADX61116.1** | 4 | Kilifi KIL36 | **PF3D7_0412400** |
| 26.6_13 | 26 | E12 | 8/43 (19%) | **ADX61577.1** | 5 | Pikine PIK06 | **PF3D7_0808600** |
| 728.4_14 | 728 | F4 | 6/39 (15%) | AAR32006.1 | 4 | PNG | PF3D7_0809100 |
| 728.4_14 | 728 | E6 | 4/39 (10%) | AGP75889.1 | 4 | Gabon MOA | PF3D7_0412700 |
| Uni1_12 | Unique | A4 | 9/38 (24%) | ACT21622.1 | 1 | Gabon 127_31 | PF3D7_1300300 |
| Uni1_12 | Unique | B1 | 5/38 (13%) | ABC95899.1 | 4 | Mali | PF3D7_0223500 |
| Uni1_12 | Unique | A3 | 4/38 (11%) | AAR31980.1 | 1 | PNG | PF3D7_0937600 |
| Uni4_14 | Unique | E10 | 5/41 (12%) | KOB63545.1 | 4 | HB3 | PF3D7_0324900 |
| Uni4_14 | Unique | E1 | 4/41 (10%) | AHI95176.1 | 3 | Malawi | PF3D7_1100200 |

**FIG. 2.**
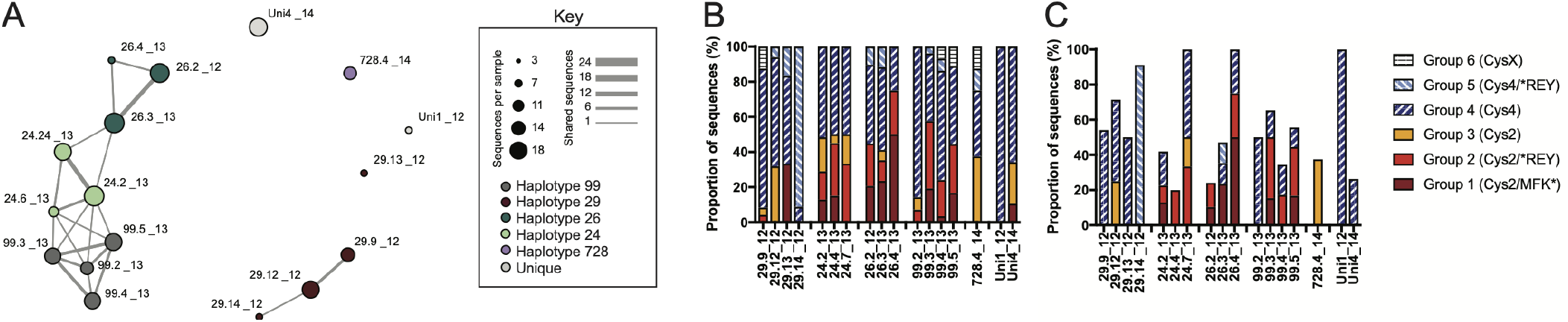
Network analysis of ex vivo DBL1*α var* sequences from CGS clustered parasites reveal relatedness within clusters. A. *var* repertoire overlap network analysis for the expressed *var* transcripts at the amino acid level. Each vertex in the diagram represents a unique parasite strain, with larger vertices indicating the total number of unique sequences in the *var* repertoire, and edge widths corresponding to the number of shared sequences between vertices (parasite isolates). Vertices have been color coded according to cluster Haplotype. B. *var* transcripts classified by cysteine/PoLV groups [13, 14] as proportion of all sequences. C. *var* transcripts classified by cysteine/PoLV groups Groups [13, 14] as proportion of dominant three sequences.

The similarity of ex vivo *var* transcription within CGS clusters can also be viewed through the lens of established *var* groupings. Studies of *var* DBL1*α* sequences have shown that they can be divided, based on sequence content, into six functionally relevant groups called cys/PoLV groups [13, 14]. We therefore also classified all transcribed *var* genes according to the six cys/PoLV groups, and plotted per-parasite *var* transcription profiles for the total expressed *var* repertoire (Figure 2B) and for the dominant three transcribed vars (Figure 2C). These results further reinforce the finding that *var* transcription is generally conserved, though not identical, by cluster.

Measurements of *var* Ups class transcript expression, by cluster, showed a similar pattern. We found that parasites within a cluster express similar Ups *var* classes ex vivo, and these *var* classes are specific to the particular cluster (Figure S3). We analyzed Ups transcription first as a ∆CT value relative to seryl tRNA synthetase, and second as a fold-change difference relative to Uni1 12—a randomly selected control strain with a unique barcode haplotype that has never been observed in another season or location among the study samples (Figure S3). While *var* Ups type transcription was conserved within clusters, each unique control strain expressed a different dominant *var* (Figure S3).

### CGS parasites express similar *var* DBL1*α* sequences and *var* Ups classes across transmission seasons

The ex vivo similarity of *var* transcription within clusters, described above, focused on clusters of parasites collected in a single year, leaving open the possibility that a single-year window might not be sufficient to observe changes of *var* transcription within CGS clusters. To better understand the longevity of these patterns, we therefore adapted parasites in short-term culture from four different CGS clusters with different persisting durations, the longest of which spanned seven years. We then asked whether transcription of *var* types measured by DBL1*α* sequence tag identity is similar across multiple years as well as within a single season (Table II; Figure 3). The caveat of such experiments is that *var* gene transcription may switch at genotype-variable rates when isolates are adapted to in vitro culture [20]. Nevertheless, we found significant overlap among DBL1*α* sequences expressed within a cluster, shown clearly in repertoire overlap networks (Figure 3A). Interestingly, Haplotype 728 showed weak but persistent overlap of transcribed sequences with Haplotype 29. To complement this analysis, we once more characterized the *var* DBL1*α* sequences using the six established cys/PoLV groups for the total expressed *var* repertoire (Figure 3B) as well as for the dominant three transcribed vars (Figure 3C) for the culture-adapted parasites, and found these cys/PoLV groups conserved by cluster as well.

**FIG. 3.**
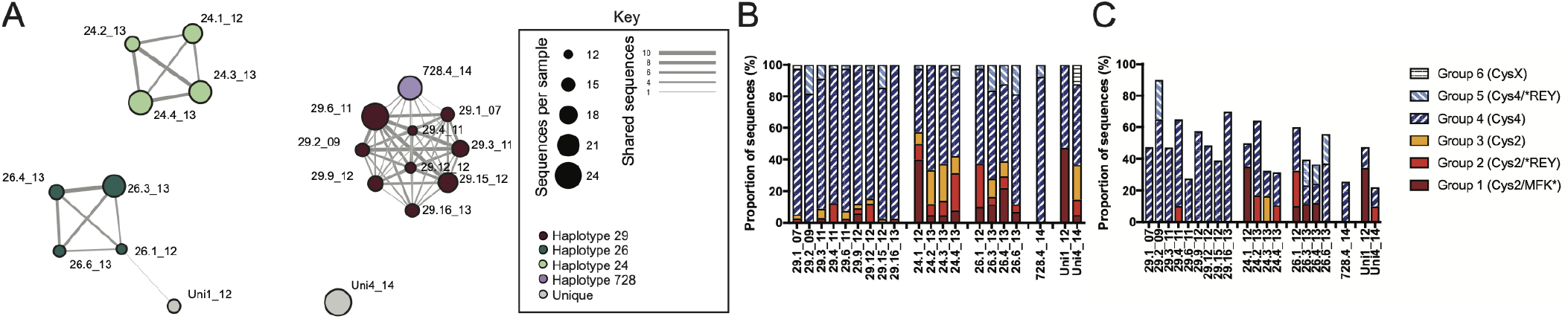
In vitro adapted CGS parasites with similar VSA recognition patterns express similar DBL1*α var* sequences. A. *var* network analysis for the expressed *var* transcripts at the amino acid level for short term adapted parasites used in VSA assays. Each vertex in the diagram represents a unique parasite strain, with larger vertices indicating the total number of unique sequences in the *var* repertoire, and edge widths corresponding to the number of shared sequences between vertices (parasite isolates). Vertices have been color coded according to cluster Haplotype. B. Var transcripts classified by cysteine/PoLV groups [13, 14] as proportion of all sequences. C. Var transcripts classified by cysteine/PoLV groups [13, 14] as proportion of dominant three sequences.

Once more, we observed a similar trend when examining Ups *var* expression patterns within clusters, even in the cluster spanning seven years (Haplotype 29, Figure S5). While occasionally an outlier was observed (for example, one parasite in Haplotype 29 strongly up-regulated UpsA3 whereas the others were not significantly different from the unique control, Figure S5), on the whole, all the parasites within the cluster exhibited similar transcriptional profiles. Also interesting was the tendency for some clusters to switch away from UpsA *var* genes and toward UpsC *var* genes, as has been described previously in culture-adapted parasites from non-immune travelers [20].

### CGS clusters show stable genomic *var* sequence diversity but not diversification over time

Identical barcodes, identical CSP, TRAP and SERA2, and similar *var* transcription hint strongly that within Senegalese CGS clusters, genomic *var* repertoires are likely to be highly similar. We confirmed that this was the case by sequencing genomic DBL1*α* domains at high depth and analyzing the resulting repertoires using multiple complementary approaches.

First, we established that the *var* repertoires within CGS clusters show a typical level of repertoire diversity by using a network approach [21] to map the *var* genes of CGS parasites onto a background network of global *var* diversity [22]. We did so both at the level of CGS Haplotypes (Figure 4A) and using individual samples within those haplotypes (Figure 4B, C, D). Reinforcing previous findings [12], we found that the CGS vertices did not cluster together in a single community but were spread throughout the network, indicating within-cluster diversity on the same order as total global diversity.

**FIG. 4.**
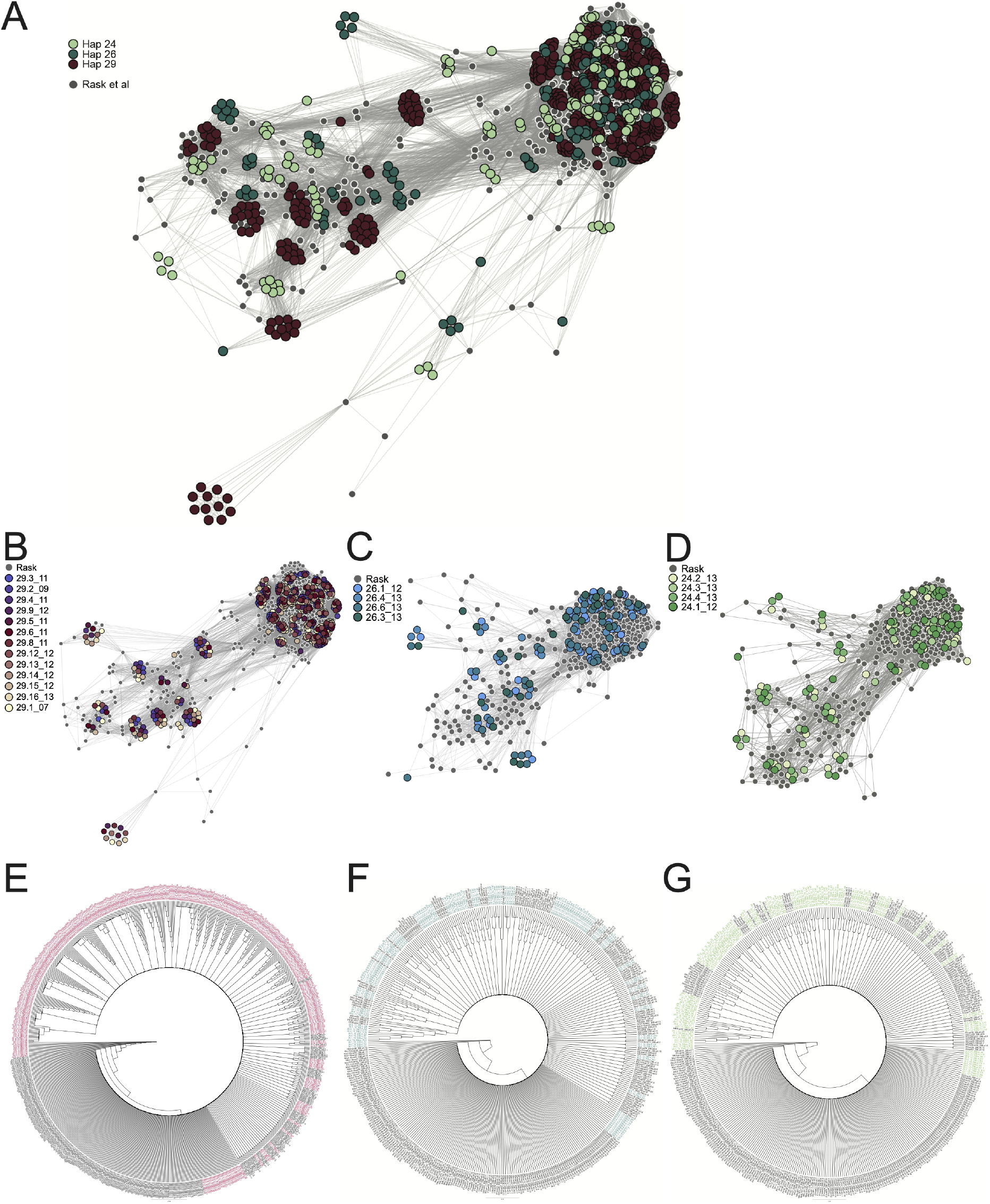
CGS parasites display conserved *var* genomic sequence diversity compared to global isolates. *var* network analysis for the expressed *var* transcripts at the amino acid level for short term adapted parasites used in VSA assays using the method described previously [21], overlaying the Senegalese isolates with sequences from 7 genomes representing global *var* diversity [22]. Each vertex in the diagram represents a unique parasite strain, and edge widths corresponding to the number of shared sequences between vertices (parasite isolates). A. Vertices have been color coded according to cluster Haplotype. B. Vertices have been color coded according to parasite isolate ID within cluster Haplotype 29. C. Vertices have been color coded according to parasite isolate ID within cluster Haplotype 24. D. Vertices have been color coded according to parasite isolate ID within cluster Haplotype 26. E-G. Neighbor-joining Phylogenetic trees using the Jukes-Cantor method were created using Geneious (version 11.0.2, created by Biomatters). Trees represent a consensus of 1000 bootstrapped replicates, showing nodes present in at least 70% of the replicate trees. Haplotype 29 (E), Haplotype 24 (F), and Haplotype 26 (G) sequences, color coded according to cluster, are aligned with global sequences from 7 parasite genomes [22], shown in black.

Next, we used a phylogenetic analysis to show that in spite of this level of *var* repertoire diversity, CGS parasites nevertheless show strong conservation of DBL1*α* sequences within Haplotype clusters, compared to the global diversity of 7 genomes (Figure 4E-G). This within-cluster conservation was most notable for Haplotype 29 (Figure 4E).

Finally, we quantified the number of *var* types that are shared between each pair of parasites by computing repertoire similarity using Bayesian repertoire overlap, a method that robustly estimates similarity in spite of incomplete repertoire coverage [19]. This analysis shows clearly that any two parasites of the same CGS Haplotype have highly overlapping *var* repertoires, and that two parasites of different Haplotypes are only minimally overlapping, near the level of expectation for parasites drawn from the global population (Figure 5). These results hold when alternative methods are used to quantify repertoire overlap (Figure S8).

**FIG. 5.**
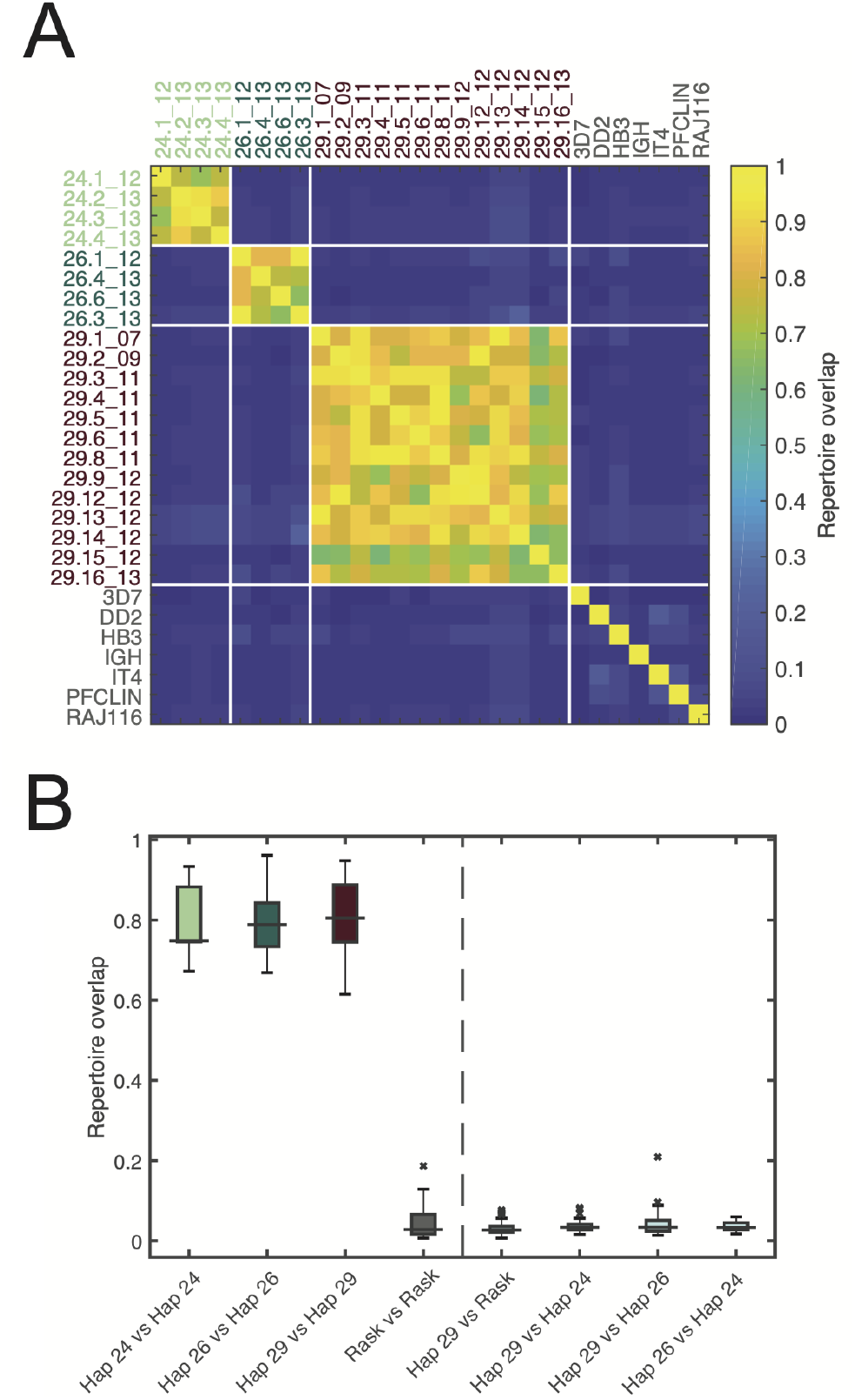
*var* repertoire overlap of CGS parasites. Overlap between *var* repertoires among CGS cluster haplotypes 29, 26, and 24, and global sequences (“Rask”), estimated via Bayesian repertoire overlap [19], are shown in two ways. A. Heatmap of all-to-all repertoire overlap estimates. B. Boxplots of repertoire overlap distributions within CGS cluster Haplotypes 29, 26, and 24 compared within cluster, between clusters, and to global sequences (“Rask”). Boxes cover interquartile ranges; lines indicate medians; whiskers extend up to 1.5×IQR. Identically constructed plots, using the pairwise type sharing method [23] are shown in Figure S8.

The genomic similarity of *var* repertoires within clusters leaves open the possibility that those repertoires are nevertheless changing over time. A recent theoretical study by Pilosof et al. [24] showed via simulations that under higher transmission levels and selection, *var* repertoires can persist over many seasons even while individual *var* genes are removed from or added to the temporally coherent repertoire. We investigated the temporal dynamics of Haplotype 29, a cluster persisting from 2007 to 2013 for which we have genomic *var* repertoire data. We observed very few unique *var* genes among cluster members and more frequently *var* genes shared by other cluster members, implying limited diversity within this cluster (Figure S6A). Nevertheless, while uncommon, we observed some evidence of SNP-level diversity as well as recombination events (Figure S6). These changes, however, were not systematic or cumulative over time. We saw no evidence that repertoires between adjacent years were more or less similar than non-adjacent years (Figure S7B), and there was no significant trend of repertoire overlap over time (Figure S7C). This result, namely that there was no significant diversification of the longstanding Haplotype 29 *var* repertoire over time, suggests that the dynamics of transmission in Senegal resulting in antigenically stable CGS clusters are unlike those used by Pilosof et al [24]. Haplotype 29 appears to have a relatively stable and not diversifying *var* repertoire genotype over seven years.

### Parasites within haplotype clusters are recognized similarly by VSA-reactive IgG

The analyses above paint a picture of Senegalese CGS haplotype clusters that are genomically and transcriptionally similar within clusters but not between, suggesting that the same may be true for recognition of surface antigens by host immunity. To test whether CGS parasites are recognized by the immune system as identical, we employed VSA recognition assays using detection by flow cytometry. Previously, both growth inhibition assays (GIA)—measuring invasion and growth inhibitory antibodies—and VSA assays—measuring antibodies that recognize parasite proteins on the infected erythrocyte surface—showed significant correlations in recognizing CGS parasites within a cluster [12]. Here we focused on VSA recognition because our preliminary data indicated that patterns of VSA recognition against CGS parasites varied by year, which have the potential to implicate this immune mechanism in selection of particular parasite genotypes over time (Figure 1).

We performed VSA assays on 18 parasite isolates representing the four CGS clusters analyzed throughout this study, with two additional parasite isolates representing unique controls from the population. The plasma samples used for recognition were selected from four consecutive years (2011-2014) in which we saw dramatic fluctuations in cluster genotype frequencies (Figure 1B), and were then chosen uniformly at random from within seasons. The resulting patterns of IgG percent recognition show strong clustering by both PCA (Figure 6A) and hierarchical clustering (Figure 6B). We confirmed this using pairwise comparisons of parasites within cluster and relative to other clusters and controls (Figure S9). Interestingly, the one exception was parasite strain 728.4 14, a member of Haplotype 728 that showed sequence identity with Haplotype 29 at both TRAP and SERA2 (Figure S2B & C) and *var* DBL1*α* transcriptional identity (Figure 3A). This strain showed high VSA correlation with parasites from the Haplotype 29 (Figure 6; S9 Figure B), and perhaps indicates some level of genome similarity between the two clusters, potentially arising from identity by descent. The results of this analysis were robust when using a rank-order approach to analyze pairwise using VSA recognition as a continuous variable, as well as when using 75th percentile cut-offs (samples with percent recognition values greater than 75% of all sample values, and thus, “highly positive”) to analyze recognition as binary “positive” or “negative” (data not shown).

**FIG. 6.**
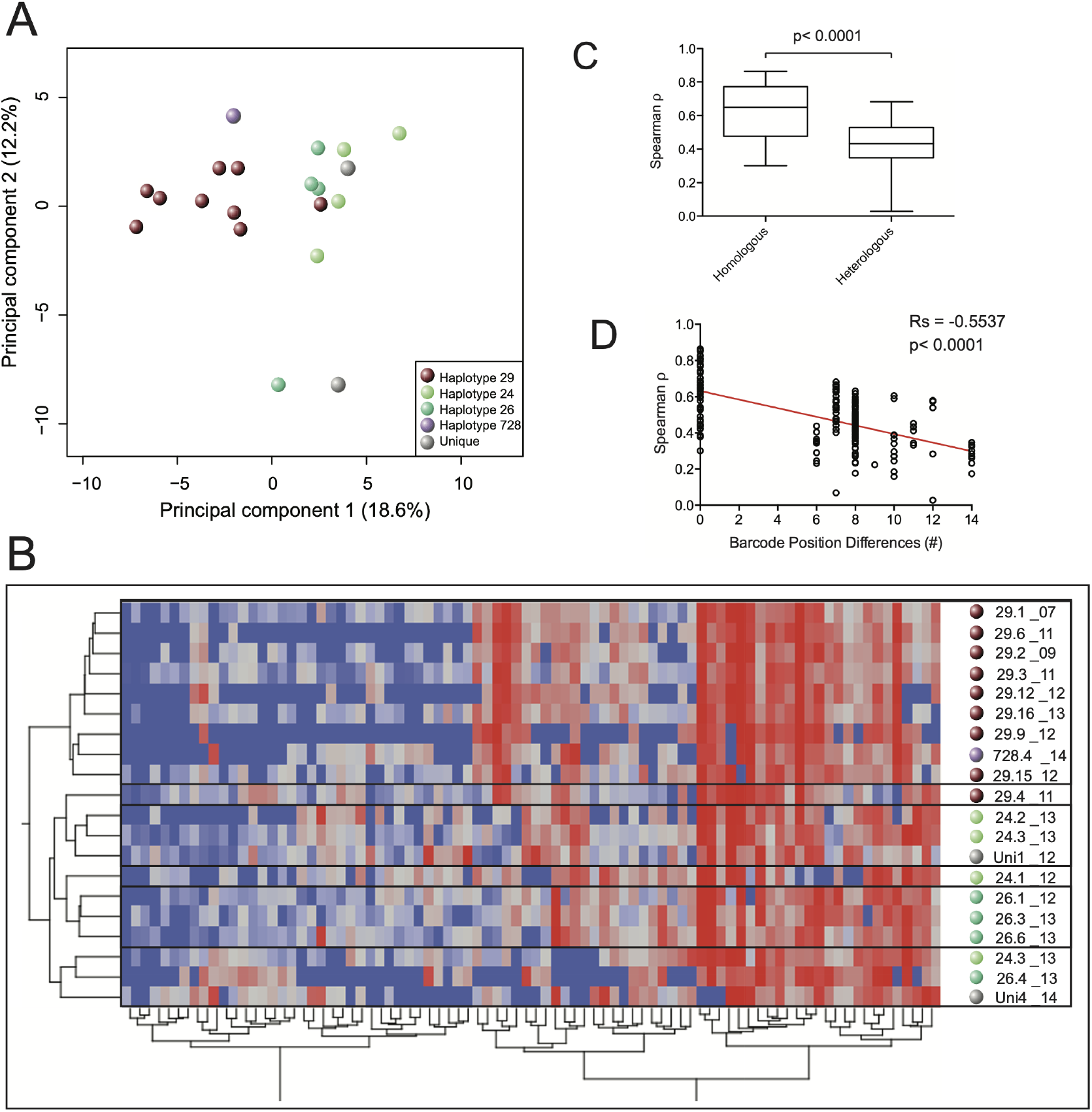
In vitro adapted CGS parasite clusters display similar patterns of immune recognition of variant surface antigens. A. PCA reveals clustering of VSA responses according to haplotype. PC1 explains 18.6% of the variance, and PC2 explains 12.2% of the variance. B. Hierarchical clustering using the Ward method shows clustering among parasites with a common haplotype. C. Overall, parasites with an identical barcode (homo) compared to a different barcode (hetero) show higher Spearman correlation coefficients in VSA response (MannWhitney test *p <* 0.0001); D. As the number of positional differences within the barcode haplotype decreases, the VSA Spearman correlation decreases as well (Linear regression: *R*^2^ = 0.3593, *p <* 0.0001). Spearman rank for Spearman correlation coefficients in VSA response and number of positional differences within the barcode is shown: *Rs* = *−*0.5537, *p <* 0.0001.

### The degree of immune recognition of CGS parasites depends on the level of the genetic similarity of the parasites

Similar patterns of VSA recognition, *var* transcription and TRAP and SERA2 sequences motivated us to determine whether the level of barcode identity (as a proxy for whole genome relatedness [3]) correlated with high Spearman rank correlation coefficients for variant surface antigen recognition determined by VSA assay. We observed a strong positive correlation between the level of barcode similarity and the VSA recognition (Figure 6C & D). This result implies that genome-level relatedness is associated with immune recognition and response at the level of variant surface antigens. The correlation between genomic similarity and VSA recognition leads naturally to the hypothesis that VSA immunity might drive selection of particular genotypes. To test this hypothesis, for parasites within each cluster, we determined the percentage of iRBC-surface positive plasma per year (those above the 75% cut-off) and compared with the cluster haplotype allelic frequencies. Unlike previous observations for Haplotype 4 [12], here we observed no significant trends between the percent positive plasma responders by year and specific haplotype frequencies (data not shown). This evidence argues against a generalizable pattern of VSA response being associated with immune selection for a particular haplotype. However, this result is based upon a limited sampling scheme, so generalizations may require further study (see Discussion).

### Multiple genotype-reactive immune responses to CGS parasites are generated at a low, but not insignificant frequency

During analysis of VSA antibody responses, we found a subset of plasma samples capable of broadly recognizing the majority of parasite tested, spanning multiple clusters. Specifically, three plasma samples among the 84 tested (3.6%) recognized more than 15 of the 20 of the parasites analyzed; and from these three, two plasma samples of the 84 tested (2.4%) recognized 19 of the 20 (95%) parasite isolates tested. These results could imply that highly multi-isolate-reactive immune responses are generated at a low, but not insignificant frequency (2.4%). Targets of these immune responses will be important areas for future studies, and further experiments would be needed to determine whether these IgG repertoires represent broadly cross-reactive responses of a few antibodies or, instead, a large repertoire of variant-specific antibodies.

## DISCUSSION

In-depth studies of P. falciparum genetic diversity often stop prior to assessing the role of such diversity on pathogenic processes, such as neutralizing immunity. Similarly, in-depth studies of virulence gene expression and pathogenic dynamics conducted with patient-isolated parasites rarely consider the role of genome-wide parasite diversity and genotypic complexity. This study sought to bridge this gap to identify the relationship of parasite genetic diversity in highly related genotypic clusters with variant gene expression and variant antigen immune recognition.

In addition to being identical by our 24-SNP molecular barcode [3], CGS parasites were also found to be highly similar at other genomic loci. Genomic identity among CGS parasites has been confirmed by the extended 96-SNP barcode, as well as whole genome sequencing [3, 25] for many of our samples. However, many highly polymorphic loci are either not included in whole genome sequence data (vars and sub-telomeric regions undergoing rapid recombination) or are potentially under strong selection and may be evolving at faster rates than the neutral SNPs used in the barcode. Therefore, we moved beyond barcode-based estimates to more deeply describe the genetic diversity of CGS clusters by sequencing highly polymorphic genetic loci—CSP, TRAP, and SERA2. We found that every parasite within a cluster contained identical CSP, TRAP, and SERA2 haplotypes and further, when these three loci were combined, each cluster mapped to its own unique haplotype that was distinct from all other clusters and controls. This finding leads us to conclude that the degree of both immune-selected diversity and recombination, even within highly polymorphic loci, is low in this population even over many years (the most dramatic example spanning 2007-2014).

The hypervariable nature of *var* genes has resulted in estimates that in highly endemic areas, the *var* repertoire is essentially unlimited [23, 26–29], with the high *R*_0_ characteristic of P. falciparum transmission being related to multiple immunogenically distinct strains with low individual *R*_0_. However, given that others have observed limited *var* gene repertoire diversity in regions of low malaria transmission [23, 30], we sought to estimate the genomic *var* diversity in our Senegalese samples. We found a dramatic lack of *var* diversity within each Haplotype cluster and conclude that isolates with identical genotypes represent essentially the same “strain” in our samples. This finding has implications for both disease control measures as well as understanding the natural dynamics of population structure and naturally acquired immunity. Further, while it is possible that *var* repertoires could be slowly diversifying over time within clusters, we found no evidence for such diversification, particularly in Haplotype 29 which spans 2007-2013.

Given the genetic identity of parasites within a CGS cluster, we investigated the transcription profiles of key virulence gene families, namely the *var* genes from both ex vivo samples from three transmission seasons (2012-2014) as well as culture-adapted parasites from clusters spanning seven transmission seasons (2007-2014). We found a striking conservation in the *var* genes expressed within clusters, both at the Ups class level as well as the sequence identity level in ex vivo samples. To delve deeper into the transcriptional relatedness of CGS parasites, we employed RNAseq to assess the level of total transcriptional variation, as well as that for key pathogenesis related gene families (e.g., invasion, *var*s, *rifin*s, and *stevor*s) across three parasite isolates within the same haplotype (Haplotype 29). Overall, two strain transcriptomes were highly similar (29.9 12 and 29.14 12) while one was distinct (29.12 12) (Figure S10A). When we compared the similarity in other variant antigen families, we observed variation depending on the specific gene family. Invasion genes were largely conserved, except for the *Clag* gene family—in which some gene family members are known to vary even within clones of the same genetic strain [31–33] (Figure S10C). *Stevor* transcription was also extremely similar for all members of the cluster, with very few significant differences observed (Figure S10D). *Rifin* gene transcription mirrored the patterns in overall transcriptome relatedness (Figure S10E). These findings imply that for the major cluster, Haplotype 29, parasites within a cluster are highly similar beyond the SNP barcode sequences, but that the degree of transcriptional similarity varies depending upon the specific variant antigen family.

While *var* gene transcription was not identical for all parasites within a cluster, each cluster nevertheless tended to express one *var* gene at a significantly higher level, a surprising finding given that “unique” parasites (isolates occurring in isolation in the population and not belonging to a cluster) each expressed a different *var* gene. While most of the ex vivo RNA samples prepared were from cluster members collected during a single season, the similar transcription profiles observed could be explained as the result of a single infected mosquito biting multiple individuals in the population. This explanation is extremely unlikely: the majority of the collection dates and locations were outside of the expected limits for both the reproductive age of the infected mosquito (assuming classic model assumptions such as 3 human bites every 10 days and an average mosquito lifespan of 10 days [34] with a maximum predicted life-span of 30 days) and mosquito flight distances (assuming a dispersal rate of 350 m/day of an uninfected mosquito) [35], and from both modeled and empirical studies finding that uninfected Anopheles mosquitoes rarely disperse beyond 3km [36, 37] (Table S2, Figure S1), with a notable recent exception [38]. Rejecting a the single-infected mosquito hypothesis implies that these samples represent parasites passaged through a mosquito at least once. Furthermore, ex vivo samples from Haplotype 26 persisted across two years (and by inference, through mosquito passage), and two members of this cluster expressed the same *var* gene (26.2 12 and 26.3 13; PF3D7 0400400). Thus, the conservation of *var* transcription ex vivo, unexplainable by a single-mosquito hypothesis, indicates that different vector and human host environments are not influencing the *var* switching rates in these parasites as much as might be expected in this low transmission environment.

Our findings are in seeming opposition to data from infections in human volunteers that showed faster *var* switching rates from pre-mosquito to acute stage infection than comparable rates observed with in vitro blood passage [39]. One possible explanation stems from P. chabaudi models, in which passage through mosquito resets virulence gene expression [40], assuming that the *var* gene expressed in our samples represents the first to be expressed post-mosquito transmission. Our data also partially corroborate a recent study from infections in non-immune human volunteers in which there was a striking conservation in the *var* genes expressed post-mosquito infection from a single-strain source of sporozoites [41].

In addition to conservation of *var* transcription ex vivo within a season or spanning two seasons, we found substantial conservation in the *var* genes expressed within clusters at the Ups class level in short-term culture-adapted parasites. In the most dramatic example, Haplotype 29 exhibited conservation of *var* Ups transcription across six non-consecutive transmission seasons. While the vars expressed after short-term adaptation were not the same as those expressed in corresponding ex vivo samples, the switched vars were conserved within the cluster. Such conservation of *var* transcription after short-term adaptation to culture indicates that the switching rates may follow a pre-programed hierarchy as all lines within a cluster expressed similar dominant *var* classes, regardless of the year they were originally isolated, and regardless of the number of implied mosquito passages.

Given that parasites within a cluster expressed highly similar variant antigens, we wanted to assess whether immune recognition of parasites within the cluster was also similar. Previous findings indicated that there is a high correlation of both GIA recognition and VSA recognition of short-term adapted parasites within the Haplotype 4 cluster [12]. Here, with short-term adapted parasites spanning seven transmission seasons, we observed a similar conservation of immune recognition of VSA antigens by cluster (Figure 6, Figure S10). This conserved recognition was most striking for Haplotype 29 where recognition profiles for eight of the nine members clustered strongly together with highly significant correlation coefficients. While Haplotypes 24 and 26 both showed significant correlation, the coefficients were moderate. Interestingly, the more distinct the parasites were by barcode, the more divergent their VSA responses were, indicating a direct relationship between the level of genomic identity and the degree of immune recognition of the particular parasite strain.

The correlation between genomic similarity and immune recognition prompted the investigation of whether there is an association with genotype frequency and immune recognition across transmission seasons for persisting clusters, which may imply immune selection of specific genotypes. Previously, we observed that for short-term adapted parasites from Haplotype 4 and individual plasmas from years spanning the emergence and decline of the cluster, there was an inverse correlation between the VSA recognition and the frequency of the haplotype in the population [12]. This interaction between immune selection and *var* diversity has been modeled to try and understand the dynamics by which populations are structured into strains or “modules” (groups of parasite genomes). Over time, modules of parasite genomes (which we call clusters), whose *var* gene combinations are more similar within a module than between modules, persist at varying rates depending on transmission and immune pressure [24]. In general, for medium and high transmission settings, models show that immune selection results in the persistence of gene combinations over longer time scales than expected under neutrality. In contrast, in low transmission settings, modular replacement was more common because immunity to genetically conserved and highly abundant modules occurs quickly [24] and selects against them, as we proposed from our previous findings [12]. In contracts, we observed the opposite: a pattern of long-term persistence only in periods of low transmission intensity. Perhaps the difference is due to the slow development of immunity in our extremely low-transmission setting which allows clusters (modules) to persist for longer periods before neutralizing immunity can clear them from the population, giving rise to new clusters. While we observed some trends with plasma spanning four of seven seasons, we observed no significant patterns of VSA recognition related to haplotype frequency. This finding does not directly rule out the possibility of immune selection of haplotypes in the population, but it argues against a unified and generalizable mechanism for all clusters. Our previous finding may have been strongly significant due to the dynamics of rapid emergence and decline of Haplotype 4, whereas for other haplotypes with dynamics of more gradual growth and decline, the relationship between immune recognition and frequency is less pronounced.

From VSA data, we also addressed the level of genotype-specific versus multiple genotype-reactive responses within the population. The level of cross-genotype recognition we observed is similar to what has been reported in a hyperendemic region of Ghana [42], measured by flow cytometry and dramatically higher than what has been previously described for VSA responses in Kenyan isolates [13], measured by agglutination. In Ghana, the authors observed frequent broad isolate recognition of up to 30-40% of 68 parasite isolates. VSA recognition was broadest against parasites causing severe disease in patients compared to mild disease. Further, the broadly recognized parasites were those isolated from younger children, indicating that the parasites infecting these children expressed common, well-recognized VSAs compared to adults with more malaria exposure [42]. In Kenya, the maximum number of parasite isolates recognized by any one child’s acute plasma was 50% (3 of 6 parasite isolates). Further, the frequency of plasma that recognized 50% of the parasite isolates (2 of 12, 16.6%) was lower than in our study (22.6%). Another study of Kenyan isolates demonstrated with a large number of adult, exposed plasma (n=557), that 3 plasma were able to react with a high number of parasite isolates (6-8 Kenyan isolates; 75-100%), meaning that 0.5% of the population had broadly reactive antibodies to VSAs [43], compared to 3.6% in our study. While the number of individuals that had broadly reactive antibodies was lower in Kenya than in our study, it is tempting to speculate that this could be due to higher population level diversity and greater effective parasite population sizes in Kenya than in Senegal. To address whether these multiple-isolate reactive IgGs represent cross-reactive responses of a few antibodies or large repertoires of variant specific antibodies, the authors cloned out and expressed specific VDJ recombination sites and tested the resulting expressed antibodies [43]. Of note, we do not observe a strong increase in the strength of responses (rank-order) by age of the plasma donors as has been observed by others in highly endemic settings [42], most likely due to the extremely low transmission intensity of the site and the equivalent risk of children and older individuals (all are equally non-immune). In order to make any solid conclusions from comparing these studies, it would be necessary to directly compare both the genetic diversity of the circulating isolates as well as the immune recognition dynamics for larger numbers of isolates over time. Further experiments defining the immune response within individuals longitudinally could shed more light on the development of neutralizing genotype-specific, or genotype-transcendent immunity to CGS clusters.

In the battle between the antigenically variant malaria parasite and the host immune system, the parasite must achieve a balance between switching too rapidly to avoid the immune response and thus exhausting the available variant antigen repertoire, and conserving the repertoire but risking clearance by the immune system [44, 45]. Few studies with clinical isolates either in vivo or ex vivo have specifically addressed *var* gene switching rates [46–49]. However, switching rates from in vitro cultures tend to be reported around 2% [50–53] whereas experimental infections in non-immune volunteers showed much higher switching rates, on the order of 18% [39, 54]. Questions have been raised by others as to whether *var* expression patterns and switching rates are driven by interactions with the environment (such as the current host’s PfEMP-1 immune responses) or whether they are hard-wired into each parasite’s genome [55]. There is likely a balance between the two, which perhaps can change depending on the level of individual and population-based immunity. It is therefore conceivable that when the level of immunity in the individual and/or population is low, there may be less of an advantage for the parasite to switch to an alternative *var* quickly [45, 56, 57], and further expression of multiple vars, rather than a single var, may not be detrimental to survival [13, 20, 56]. Such a dynamic is similar to what we observe in Senegal where parasites with the same genetic barcode express strikingly similar *var* genes over many years. Further, as the population-level parasite genetic diversity is expected to decrease, the effective parasite population size can also decrease, though genotypic richness can also decrease in the absence of a significant change in effective population size [58]. The effective population size in Thiès, Senegal was estimated to have dropped from 402 in 2006 to approximately 10 in 2011 [3], which almost certainly resulted in the number of parasite antigenic types becoming more restricted. Such restriction could result in a focused population-level immune response, giving rise to immune escapee genotypes—potentially contributing to population rebound in genotypic richness. Interestingly, in 2012-2013, precisely such a rebound was observed, with the effective population size estimated to be 38 and rising [3].

Taken together, this study represents the largest and most comprehensive analysis of individual persisting parasites from the synthesis of genetic, transcriptional, immunological, and host-pathogen interaction data, to date. Such systems data are essential if we are to uncover the mechanisms for parasite persistence in the population. Whether the immune response to specific parasite genotypes plays a role in the recent parasite genotypic richness rebound has yet to be determined. However, it is clear is that CGS parasites are genetically, transcriptionally, and antigenically extremely similar, and these similarities can be observed from host to host and even from year to year. The implications of such conservation are only now being revealed in the population, and close monitoring of both parasite diversity and immune recognition in the population and in the individual, over time, are required. Such future observations would provide valuable information on the development and potentially protective nature of immune responses to CGS parasite clusters, as well as protective immunity directed against novel target antigens, which could provide cross-protection against diverse P. falciparum isolates. Additionally, the immunological conservation among CGS types could be exploited as a tool to understand parasite population movement and dynamics of decline and rebound in the context of malaria elimination interventions.

## MATERIALS AND METHODS

### Ethics Statement

Both the Institutional Review Board of the Harvard T.H. Chan School of Public Health (CR-16330-01) and the Ethics Committee of the Ministry of Health in Senegal (0127MSAS/DPRS/CNRES) approved this study. All samples were collected after written informed consent according to the ethical requirements of both Institutional Review Boards.

### *P. falciparum* parasite isolates

We enrolled patients in the study by passive case detection for malaria-like symptoms. Patients visiting the SLAP (Service de Lutte Anti-Parasitaire) clinic in Thiès, Senegal who were confirmed positive for P. falciparum malaria by rapid diagnostic testing (when available) and microscopy slide analysis were offered the opportunity to enroll in the study. Both venous blood samples (5 ml) and finger prick blood spotted onto filter paper were collected from consenting patients. Upon transport to the laboratory in Dakar, erythrocytes and plasma were separated by centrifugation. Plasma was retained and stored at −20o C prior to VSA assay. Infected erythrocytes were frozen in glycerolyte for in vitro culture adaptation. Ex vivo short-term parasite cultures were initiated, and RNA harvested in the first generation lysed with TRI reagent BD prior to RNA extraction. In this study, we focus on seven clusters of identical (CGS) parasites. Monogenomic status of parasites in these clusters was determined by Msp1 and Msp2 typing and by 24 SNP molecular barcoding from filter paper taken directly from patients. Monogenomic parasite isolates included in this study were short-term in vitro culture adapted (no more than 15 generations) prior to in vitro immune assays. Periodic molecular barcode analysis was performed during the culture adaptation process to confirm no cross-contamination with other genotypes and the specific barcode signature of each strain. Culture adaptation success rate was 100%, following the previously described protocol [59]. Original identifiers of parasite isolates have been re-coded to reflect the barcode haplotype and the year of collection, as described in detail in Table S1.

### Parasite genotyping

Genomic DNA (gDNA) was extracted from both whole blood spotted onto Whatman FTA filter paper (Whatman) and culture-adapted parasites. DNA was extracted from filter paper punches using the manufacturer’s protocol for Promega Maxwell DNA IQ Casework Sample kit (Promega) or Qiagen DNA Blood kit (Qiagen). A molecular barcode genotype was generated for each sample, as previously described [5]. Sequencing of highly polymorphic loci (CSP C-terminus/PF3D7 0304600/CDS 858 – 1190bp; TRAP/PF3D7 1335900/CDS 1,222 – 1,581bp; and SERA2/PF3D7 0207900/CDS 72 – 357bp) [60] was performed in addition to molecular barcoding for each culture-adapted parasite strain using two rounds of PCR amplification. PCR primers (CSP-C-terminus, TRAP, SERA2; Plasmodium sequence in bold) are described in Table S3.

First-round PCR products for CSP-C-terminus and TRAP were amplified with the following reaction mixture: 2.5*µ*l 10X Platinum Taq High Fidelity Buffer, 0.5*µ*l dNTPs (10mM stock, Denville Scientific), 1*µ*l 50mM MgCl2 (Invitrogen), 0.5*µ*l forward primer (10*µ*M stock), 0.5*µ*l reverse primer (10*µ*M stock), 0.2*µ*l Platinum Taq High Fidelity Polymerase, 17.8*µ*l of Diethylpyrocarbonate (DEPC)-treated water to which 2 *µ*l of gDNA was added for each strain. First-round PCR products for SERA2 were amplified with the following reaction mixture: 2.5*µ*l 10X Platinum Taq High Fidelity Buffer, 0.5*µ*l dNTPs (10mM stock, Denville Scientific), 1*µ*l 50mM MgCl2 (Invitrogen), 1*µ*l forward primer (10*µ*M), 1*µ*l reverse primer (10*µ*M), 0.2*µ*l Platinum Taq High Fidelity Polymerase, 16.8*µ*l of DEPC water to which 2 *µ*l of gDNA was added for each strain. Thermal cycling conditions were as follows: 95^*◦*^C for 5 min, 30 cycles of [94^*◦*^C 30 sec, 60^*◦*^C 30 sec, 72^*◦*^C 1 min] and a final extension of 3 min at 72°C.

Second-round PCRs for CSP-C-terminus, TRAP and SERA2 were performed with the following reaction mixture: 2.5*µ*l 10X Platinum Taq High Fidelity Buffer, 0.5*µ*l dNTPs (10mM stock, Denville Scientific), 1*µ*l 50mM MgCl2 (Invitrogen), 1*µ*l forward primer (10*µ*M), 1*µ*l reverse primer (10*µ*M), 0.2*µ*l Platinum Taq High Fidelity Polymerase, 17.8*µ*l of DEPC water to which 1 *µ*l of first-round PCR product was added for each strain.

Thermal cycling conditions for CSP-C-terminus and TRAP were as follows: 50°C for 2 min, 70°C for 20 min, 95°C for 10 min, 5 cycles of [95°C 15 sec, 60°C 30 sec, 72°C 1 min], 1 cycle of [95°C 15 sec, 80°C 30 sec, 60°C 30 sec, 72°C 1 min], 4 cycles of [95°C 15 sec, 60°C 30 sec, 72°C 1 min], 1 cycle of [95°C 15 sec, 80°C 30 sec, 60°C 30 sec, 72°C 1 min], 4 cycles of [95°C 15 sec, 60°C 30 sec, 72°C 1 min], 5 cycles of [95°C 15 sec, 80°C 30 sec, 60°C 30 sec, 72°C 1 min]. Thermal cycling conditions for SERA2 were as follows: 95°C for 5 min, 9 cycles of [94°C 30 sec, 60°C 30 sec, 72°C 1 min] and 72°C for 3 min.

PCR products were sequenced by Sanger sequencing in both forward and reverse directions. Contiguous sequences (contigs) were generated using SeqMan (Lasergene 10). Sequence alignments were performed with MacVector (version 12.7.3), sequences were trimmed based on the P. falciparum specific portion of the primers and high sequence quality, and neighbor-joining Phylogenetic trees using the Jukes-Cantor method and the PhyML algorithm [61] were created using Geneious (version 11.0.2, created by Biomatters) and formatted using FigTree (version 1.4.3). Trees represent a consensus of 1000 bootstrapped replicates. For the CSP-TRAP-SERA2 combined haplotype, the same sequences that were used for individual haplotypes were combined for the merged haplotype, specifically 279 nucleotides for CSP, 328 nucleotides for TRAP, and 231 nucleotides for SERA2. There were 2 samples for which we did not have high quality sequence reads at 1 locus (24.7 13 for TRAP and 26.1 12 for SERA2) so these samples were excluded from the merged haplotype analysis.

### RNA and cDNA preparation

RNA from synchronous parasite cultures was prepared using initial lysis with TRI reagent BD (Molecular Research Center), followed by phenol chloroform extraction, overnight ethanol precipitation, and purification on PureLink RNA Mini columns (Invitrogen). RNA was treated with Turbo DNase (Invitrogen), and cDNA was synthesized using SuperScript III and random hexamers (Invitrogen).

### VSA binding by flow cytometry

Synchronous late-trophozoite to mid-schizont stage parasite cultures were diluted to 1% hematocrit to prevent clumping and washed twice with 1 × PBS (phosphate buffered saline) + 2% FBS (fetal bovine serum). IgG surface staining was assessed as previously described [62] with minor modifications in the dye combinations used. Here, we use 1*µ*g/ml final concentration of ethidium bromide to stain parasitized erythrocytes and 4 *µ*l/ml Alexa Fluor 488-conjugated goat anti-human IgG (H+L) (Invitrogen, Life Technologies), based on optimal signal to noise between positive control plasma (10 Mali highly exposed individual plasma) and negative control plasma (10 Boston unexposed individual plasma). Test plasma samples were randomly selected from 2011 (n=21), 2012 (n=21), 2013 (n=21) and 2014 (n=21), based on available sample volume. Plasma samples were subjected to a specificity criterion of having background IgG reactivity of no greater than 5% (in general, the maximum observed background staining of unexposed plasma) for two different donors of uninfected erythrocytes. Any sample that failed to meet this criterion was excluded from analysis. Of the n=92 plasma samples tested (n=84 samples from Senegal and n=8 unexposed Boston controls), all met this criterion, thus all plasmas were included in the final analysis. In addition to measuring percent IgG positive (%), we scored plasma samples as responders or non-responders based on a 2*σ* cutoff (mean + two standard deviations of eight unexposed Boston plasmas). Any sample that was negative by cut-off received a rank of zero and all other positive plasma were ranked in ascending order. We further applied a quartile-based cut-off (Q3, 75th percentile) to select the very highly positive responders by strain. This allowed us to compare “percent positive responders” across all experiments. Additionally, we calculated the “VSA antibody repertoire score” [13]—a score that determines the number of parasite genotypes recognized by an individual plasma. Positive recognition was determined using a 75th percentile cut-off, meaning that the individual sample had a higher percent recognition than 75% of the individual plasma samples tested.

### Principal components analysis of VSA response

We performed principal components analysis (PCA) on the VSA responses for each parasite using the Row-wise method (JMP 12.0), to search for structural patterns among haplotypes. Each parasite strain is represented as a point on a plane defined by pairwise functions of the three first principal components (PC1 and PC2, and by PC2 and PC3).

### *var* gene Ups class transcriptional profiling

Quantitative PCR was performed on cDNAs using iTaq Universal SYBR Supermix (BioRad) with an ABI Prism real time PCR machine. Primers used in these experiments have been previously described [12, 63]. Briefly, cDNA (RT+ and RT-) was diluted 1/10 with DEPC grade water and 1*µ*l was used per reaction. The reaction mixture was as follows for each primer pair: 1*µ*l cDNA, 10*µ*l SYBR Green, 4*µ*l primer mix (equal parts of forward and reverse at 2.5*µ*M; 0.5*µ*M final concentration), and 5*µ*l DEPC grade water. Samples were run in triplicate with the exception of controls seryl tRNA synthetase and fructose biphosphate aldolase, which were run in sextuplicate. Reaction conditions were as follows: 95°C for 10:00 (initial denaturation), followed by [95°C for 0:15, 54°C for 0:40, 60°C for 1:00], repeated for 40 cycles with no final extension.

Data were analyzed as ∆CT relative to seryl tRNA synthetase. Fold change was calculated as 2^*−*∆∆CT^ compared to one of the unique controls (Uni1 12).

### *var* gene DBL1*α* sequencing

Three to four randomly selected parasite isolates from clusters of interest (Haplotypes 24, 26, 29, 99, 728, and unique) were used to generate *var* DBL1*α* sequences. DBL1*α* regions were amplified from cDNAs by PCR as previously described [64]. Here, we focused on one set of primer pairs to amplify sequence tags of DBL1*α*: Primer pair 2: *α*-AF/*α*BR (375-475bp products) [65]. All PCR products were amplified with the following reaction mixture: 2.5*µ*l 10X Platinum Taq High Fidelity Buffer, 0.5*µ*l dTNPs (10mM stock, Denville Scientific), 1*µ*l 50mM MgCl2 (Invitrogen), 0.5*µ*l forward primer (10*µ*M), 0.5*µ*l reverse primer (10*µ*M), 0.2*µ*l Platinum Taq High Fidelity Polymerase, 18.8*µ*l of DEPC water to which 1 *µ*l of cDNA or gDNA was added for each strain. RT+ and RT-reactions were performed for all PCR pairs. PCR amplification was performed using the following program: 95°C for 5:00, 42°C for 1:00, 60°C for 1:00 followed by [94°C for 1:00, 42°C for 1:00, 60°C for 1:00], repeated for 29 cycles with no final extension [65]. PCR products were cloned into pCR 4.1 TOPO (Invitrogen), transformed into Chemically Competent E. coli XL10 Gold cells (Agilent), and plated on agar with 100*µ*g/ml ampicillin. A total of 48 colonies per PCR reaction were selected and sequenced in both the forward and reverse direction by Sanger sequencing. Only sequences that yielded good quality sequence (sequences with a Phred quality score of *Q* ≥ 30) in both directions were included in analysis.

### *var* sequence analysis

Sequence reads were analyzed individually. Contiguous sequences were generated using Geneious (version 8.1, created by Biomatters) and translated in all 6 frames. Translated consensus sequences were scanned for the RSFADIG motif described previously [64]. For sequences containing the RSFADIG motif, alignments of consensus sequences were performed using Clustal W.

### *var* transcript dominance

The dominance of *var* transcripts for each patient was scored according to the method described by Normark et. al. using the following equation: fij = rij/ni, (rij = the number of sequences for patient i in cluster j and ni = total number of reads from each patient i) for each primer pair [64]. *var* sequences identified by BLAST as PFE1640w (PF3D7 0533100 or PFIT 0536800; varCOM-MON or var1CSA)), were excluded from subsequent analysis.

### *var* DBL*α* repertoire network

Translated sequences containing both a RSFADIG motif and a PQ**R motif, and not identified as PFE1640w (PF3D7 0533100 or PFIT 0536800; varCOMMON or var1CSA) were included in *var* network analysis. For each parasite strain, nucleotide sequences were clustered and replicate sequences were counted and then condensed into a single instance. The resulting repertoires of *var* sequences were compared at the nucleotide level to count the number of shared sequences between every pair of repertoires. This formed a network in which each vertex is a strain and each edge between two isolates is labeled with the integer number of shared *var* sequences. In network diagrams, the size of each vertex represents the size of the expressed repertoire and the edge widths correspond to the edge weights, indicating the number of shared sequences between the connected pair of vertices. Vertex colors represent specific barcode haplotypes for the corresponding strain. Figures were generated using webweb software [66].

### Estimating *var*repertoire overlap

Given complete sets of sequences of two parasites’ *var* repertoires, the overlap between repertoires is calculated by counting the number of *var* sequences shared by both parasites. However, due to the absence of complete sequence data, we estimated the number of shared sequences using the unbiased method of Bayesian repertoire overlap [19], which accounts for the uncertainty of estimates due to sample size. We generated posterior distributions of repertoire overlap, assuming repertoire sizes of 40, based on the average total number of DBL1*α* sequences in the 3D7, DD2, and IT4 genomes [22]. Unbiased estimates, based on the posterior mean [19], were used in constructing the heatmap and data in Figure 5. To produce sparse graphs, Figure S4A shows maximum a posteriori estimates and Figure S4B shows the lower bound of the 95% credible interval indicating the edges in the pairwise-overlap network for which there exists overwhelming evidence of existence. For historical comparison, we also estimate *var* repertoire overlap using pairwise type sharing [23], for which heatmaps are shown in Figure S8.

### *var* DBLf*α* cysteine/PoLV groups

Var DBL1*α* sequences were categorized based on the number of cysteines and size into cysteine/PoLV groups, as defined by others [13, 14].

### RNAseq analysis

200ng of total RNA was sent to Beijing Genomics Institute (Tai Po, Hong Kong) for RNAseq data generation. Enrichment of cDNA transcripts was performed using polyA selection and raw read data were analyzed at the Broad Institute (Cambridge, MA). Reads were trimmed with Trimmomatic, ends of reads were removed if phred quality was less than 15 [67], reads were aligned to the 3D7 reference genome with TopHat [68], transcript expression was measured with CuffLinks [69] and samples were compared with CuffDiff2 [70]. Statistical analysis was performed using the fragments per kilobase per million (FPKM) metric values for each locus.

## ACKNOWLEDGEMENTS

We acknowledge all the study participants in Thiès, Senegal, for agreeing to participate in this work, as well as Aminata Mbaye, Baba Dieye, Yaye Die Ndiaye, Papa Elhadj Omar Gueye, Younous Diedhiou, Lamine Ndiaye, Amadou Mactar Mbaye, and Ngayo Sy for assistance with sample collection and processing. We acknowledge Katelyn Durfee and Caleb Irvine for assistance with molecular barcoding. We thank Selina Bopp for critical reading of the manuscript. We thank Sidiya Mbodj and Fatoumata Dabo for assistance with geolocation for Thiès.

## SUPPLEMENTAL MATERIALS

**FIG. S1.**
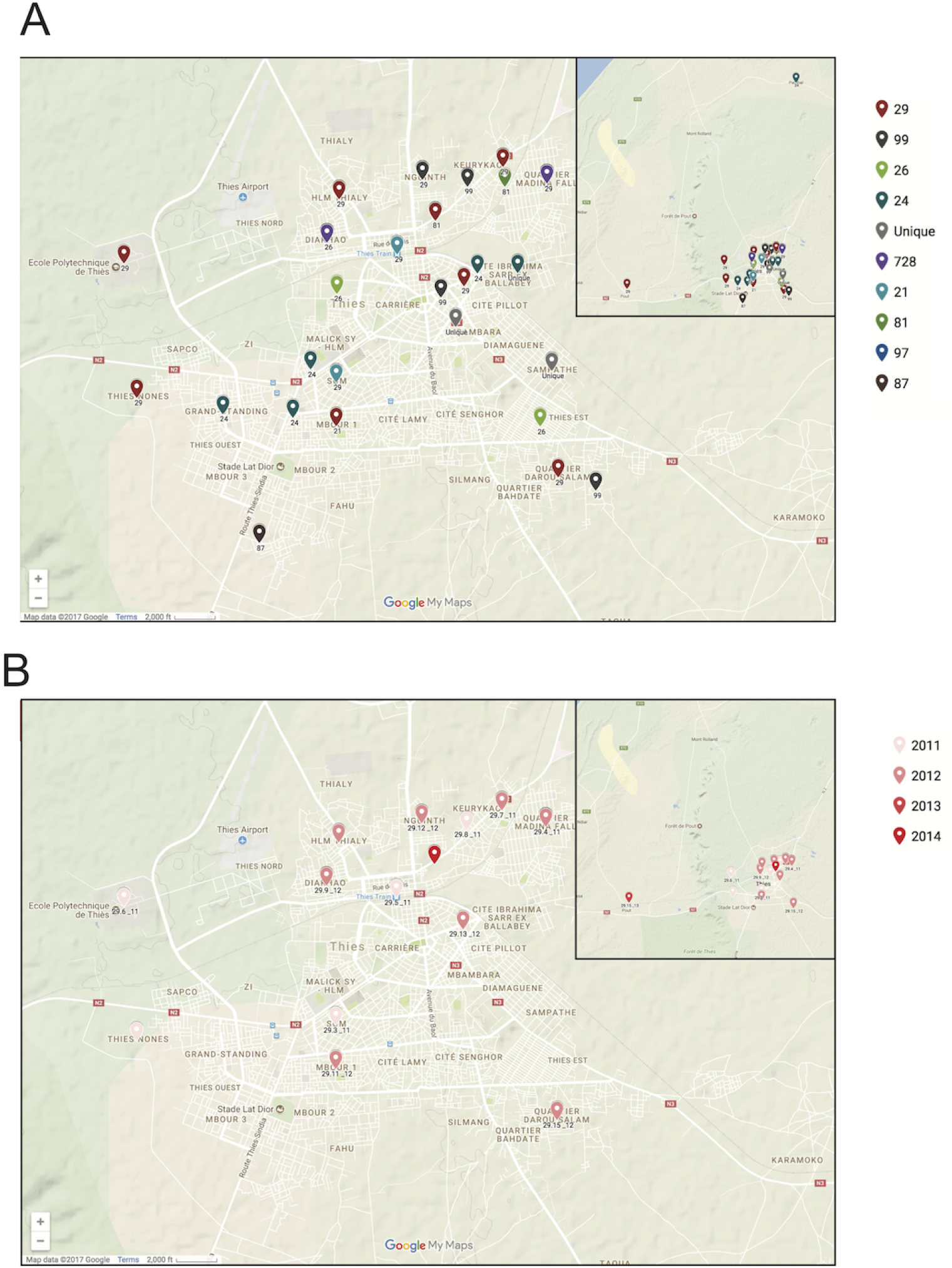
Spatio-temporal locations of CGS haplotypes. A. Spatial mapping of CGS haplotypes shown over all years, colored by haplotype. B. Spatio-temporal mapping of Haplotype 29 shown over all years, colored by year of collection and coded by sample haplotype ID.

**FIG. S2.**
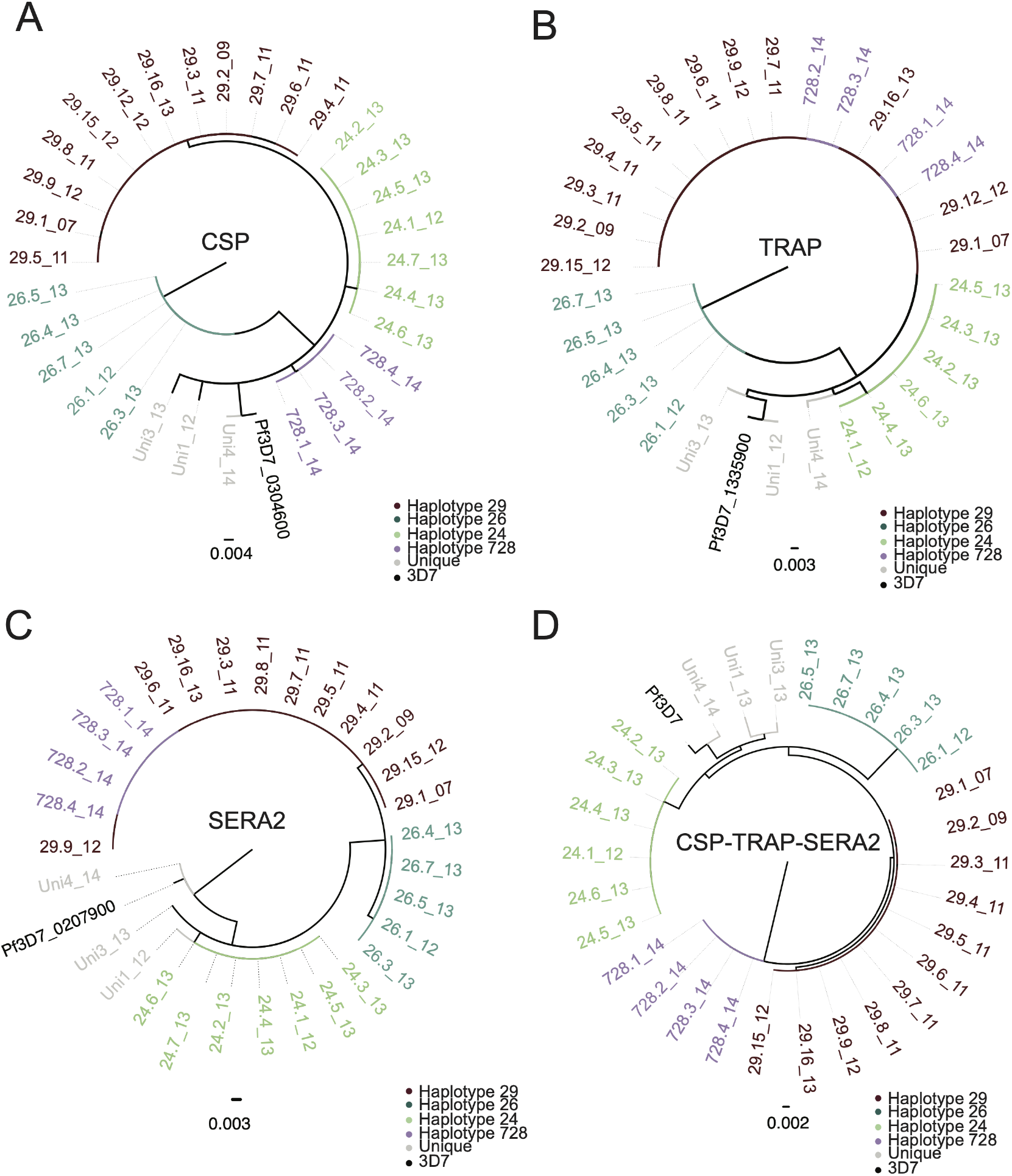
CGS genetic identity at highly polymorphic loci. CGS clusters share identical highly-polymorphic sequences for (A) CSP, (B) TRAP, and (C) SERA2. D. When CSP, TRAP and SERA2 sequences are combined into a single haplotype, parasites within CGS clusters share identical haplotypes, which are completely unique from other clusters.

**FIG. S3.**
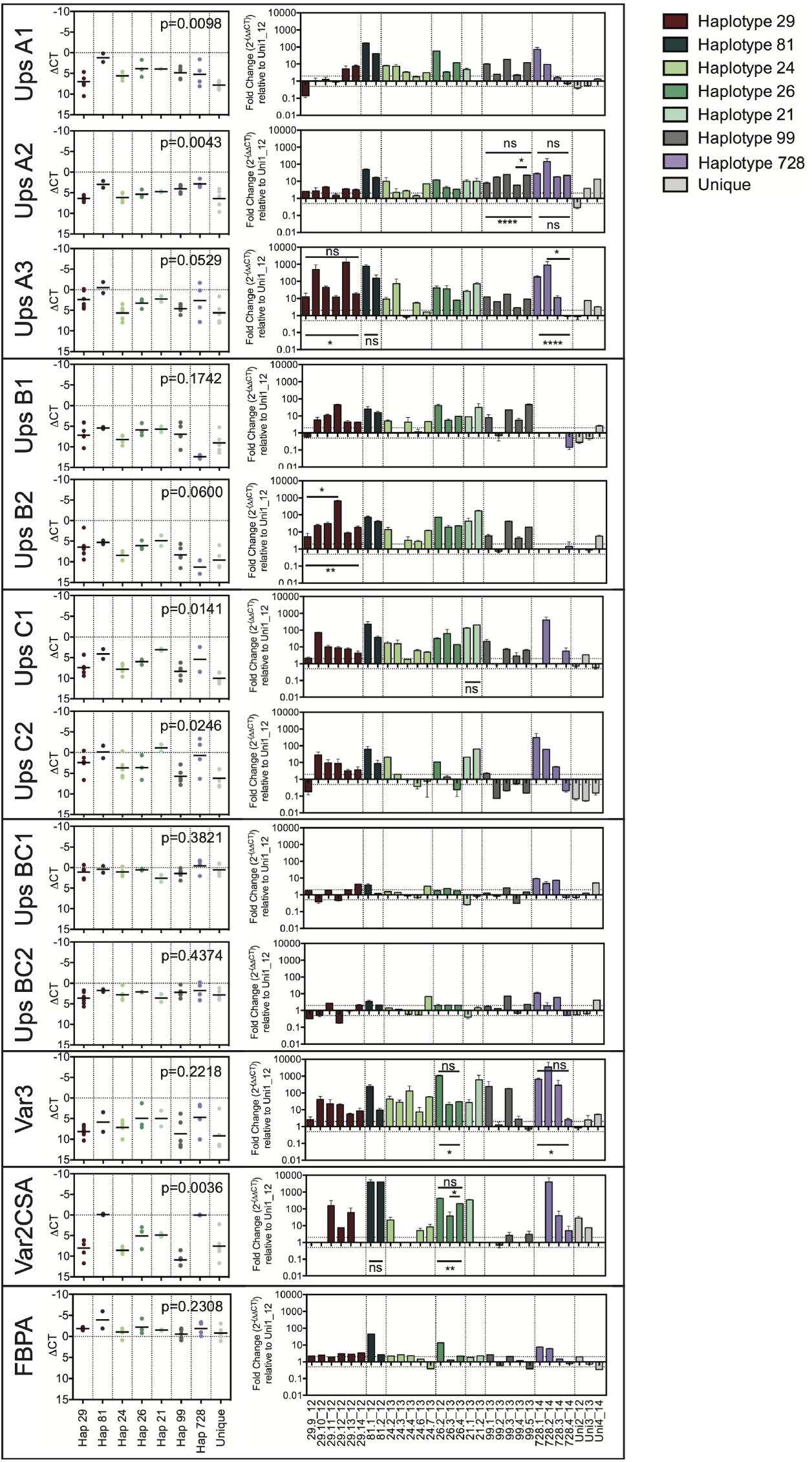
Ex vivo Ups *var* transcription in CGS clustered parasites. *Var* promoter region Ups qPCR was performed on ex vivo samples to identify the var types expressed by CGS clustered parasites and controls. Data was calculated as ∆CT relative to seryl tRNA synthetase (left) and fold change is expressed relative to the unique control strain Uni1 12 (right). Error bars for fold change represent the standard error of the mean for triplicate values. *P*-values shown on the left are from Kruskal-Wallis for *δ*CT comparisons of each cluster. Individual sample comparisons of fold change on the right are shown for the dominant var/s for each haplotype, with *p*-value significance among samples within a cluster (from Kruskal-Wallis with Dunn’s test for multiple comparisons for clusters of 3 or more; or Mann-Whitney U test for clusters with 2 samples) shown below the bars of the graph, and *p*-value significance for individual comparisons within the cluster shown above the bars. When all individual comparisons are not-significant, “ns” is shown above a bar spanning all samples and any significant outliers are marked with their *p*-value significance above a bar spanning the specific sample comparison. Significance levels are as follows: ns, *P >* 0.05; ^*\**^, *P <* 0.05; ^*\*\**^, *P <* 0.01; ^*\*\*\**^, *P <* 0.001, ^*\*\*\*\**^, *P <* 0.0001. Fructose bisphosphate aldolase (FBPA) is shown as an additional control.

**FIG. S4.**
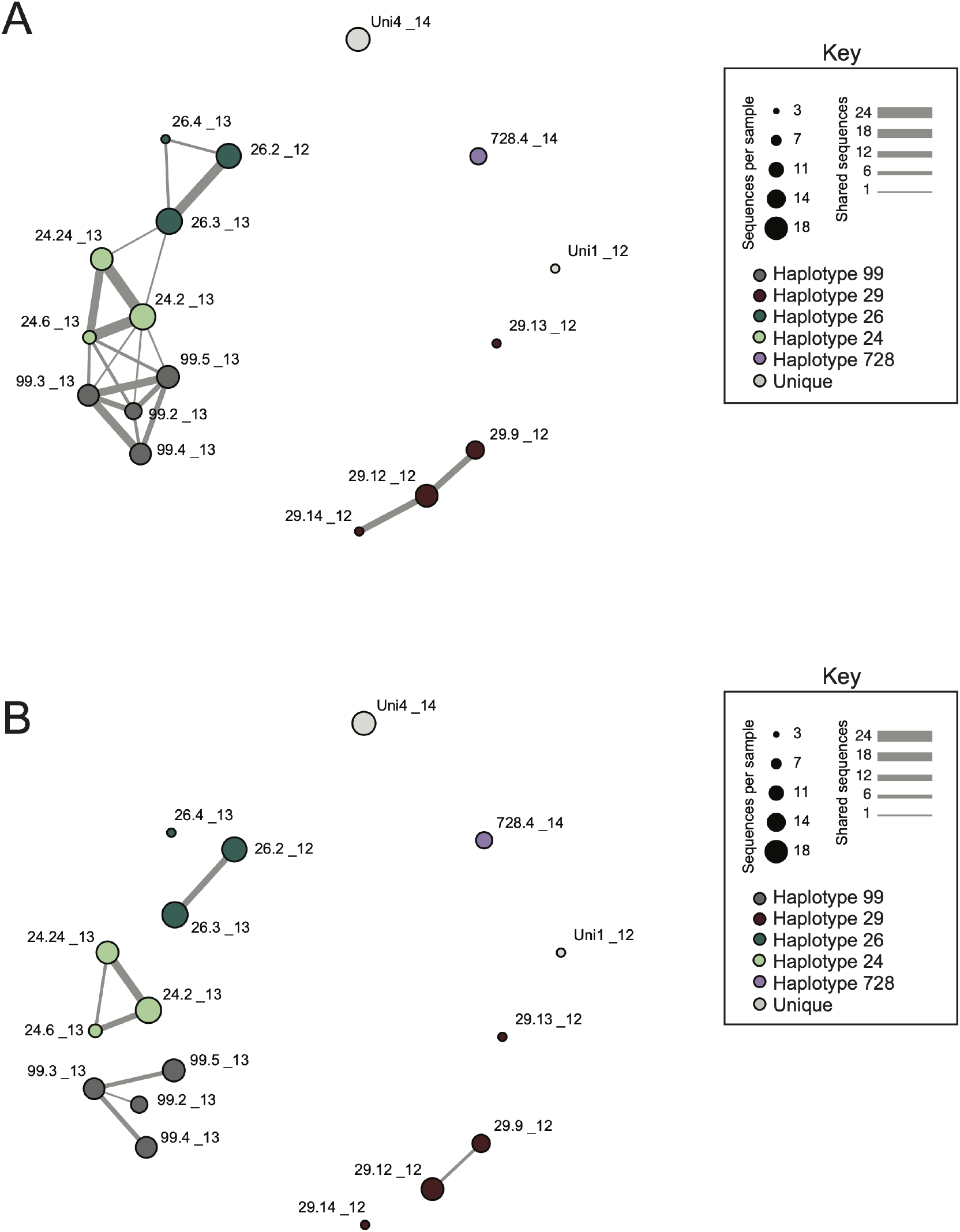
DBL1*α* var network maximum likelihood estimates. A. Confidence in the *var* repertoire overlap based on maximum likelihood estimates accounting for the uncertainty due to sample size. Each vertex in the diagram represents a unique parasite strain, with larger vertices indicating the total number of unique sequences in the var repertoire, and edge widths corresponding to the maximum likelihood estimate of the number of shared sequences between vertices (parasite isolates) assuming complete sampling (*n* = 40) of the *var* DBL repertoire. Vertices have been color coded according to cluster Haplotype. B. Confidence in the *var* repertoire overlap based on maximum likelihood estimates using the lower boundary of the 95% credible interval, representing the most stringent and conservative criteria for true pairwise-overlap.

**FIG. S5.**
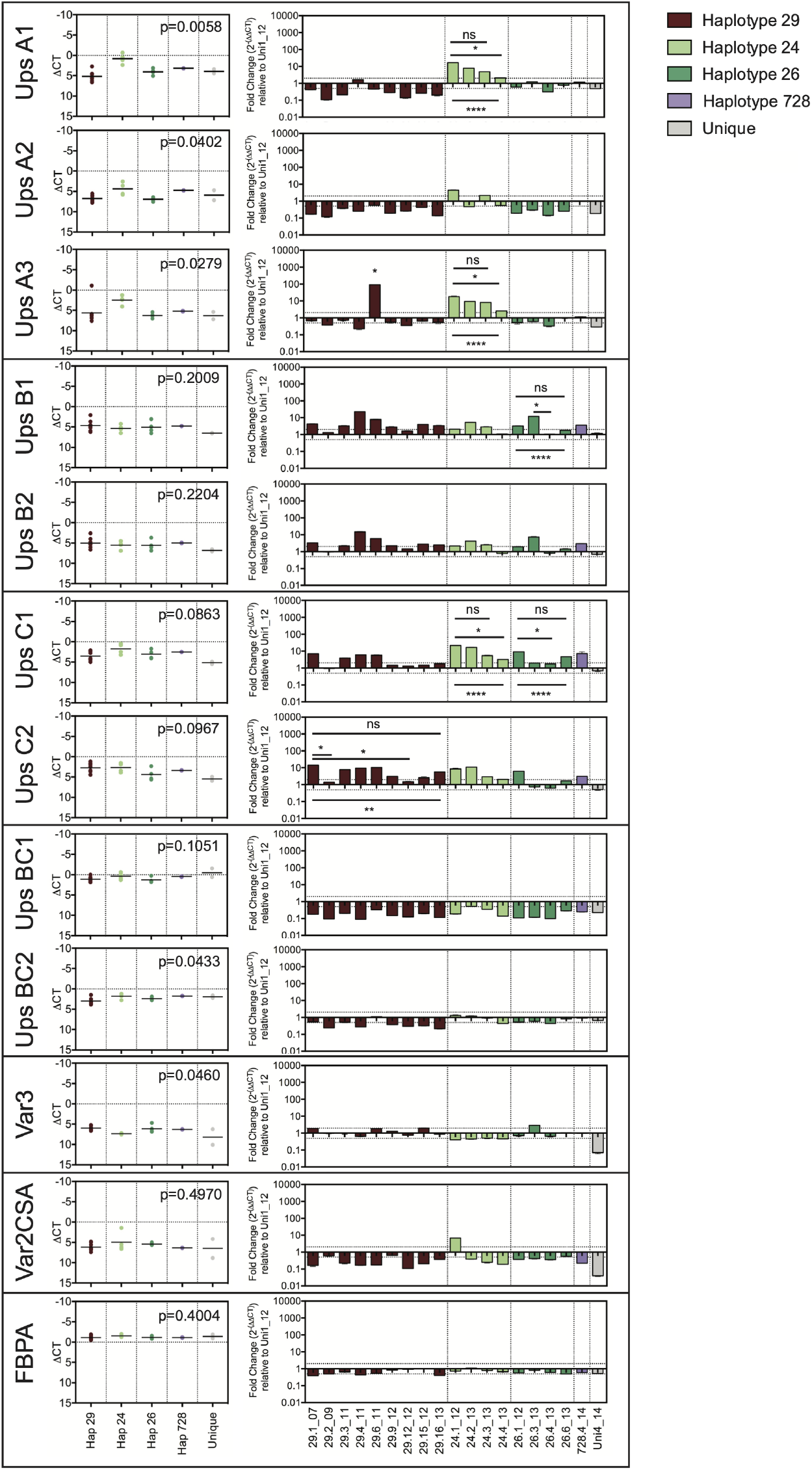
In vitro adapted CGS parasites with similar VSA recognition patterns express similar *var* Ups classes. *Var* promoter region Ups qPCR was performed on VSA matched samples (RNA isolated from the same parasite cycle used in VSA recognition assays) to identify the *var* types expressed by CGS clustered parasites and controls for which VSA phenotyping has been performed. Data was calculated as ∆CT relative to seryl tRNA synthetase (left) and fold change is expressed relative to the unique control strain Uni1 12 (right). Error bars represent the standard error of the mean for triplicate values. *P*-values shown on the left are from Kruskal-Wallis for ∆CT comparisons of each cluster. Individual sample comparisons of fold change on the right are shown for the dominant var/s for each haplotype, with p-value significance among samples within a cluster (from Kruskal-Wallis with Dunn’s test for multiple comparisons) shown below the bars of the graph, and *p*-value significance for individual comparisons within the cluster shown above the bars. When all individual comparisons are not-significant, “ns” is shown above a bar spanning all samples whereas any significant outliers are marked with their *p*-value significance above a bar spanning the specific sample comparison. Significance levels are as follows: ns, *P >* 0.05; ^*\**^, *P <* 0.05; ^*\*\**^, *P <* 0.01; ^*\*\*\**^, *P <* 0.001, ^*\*\*\*\**^, *P <* 0.0001. Fructose bisphosphate aldolase (FBPA) is shown as an additional control.

**FIG. S6.**
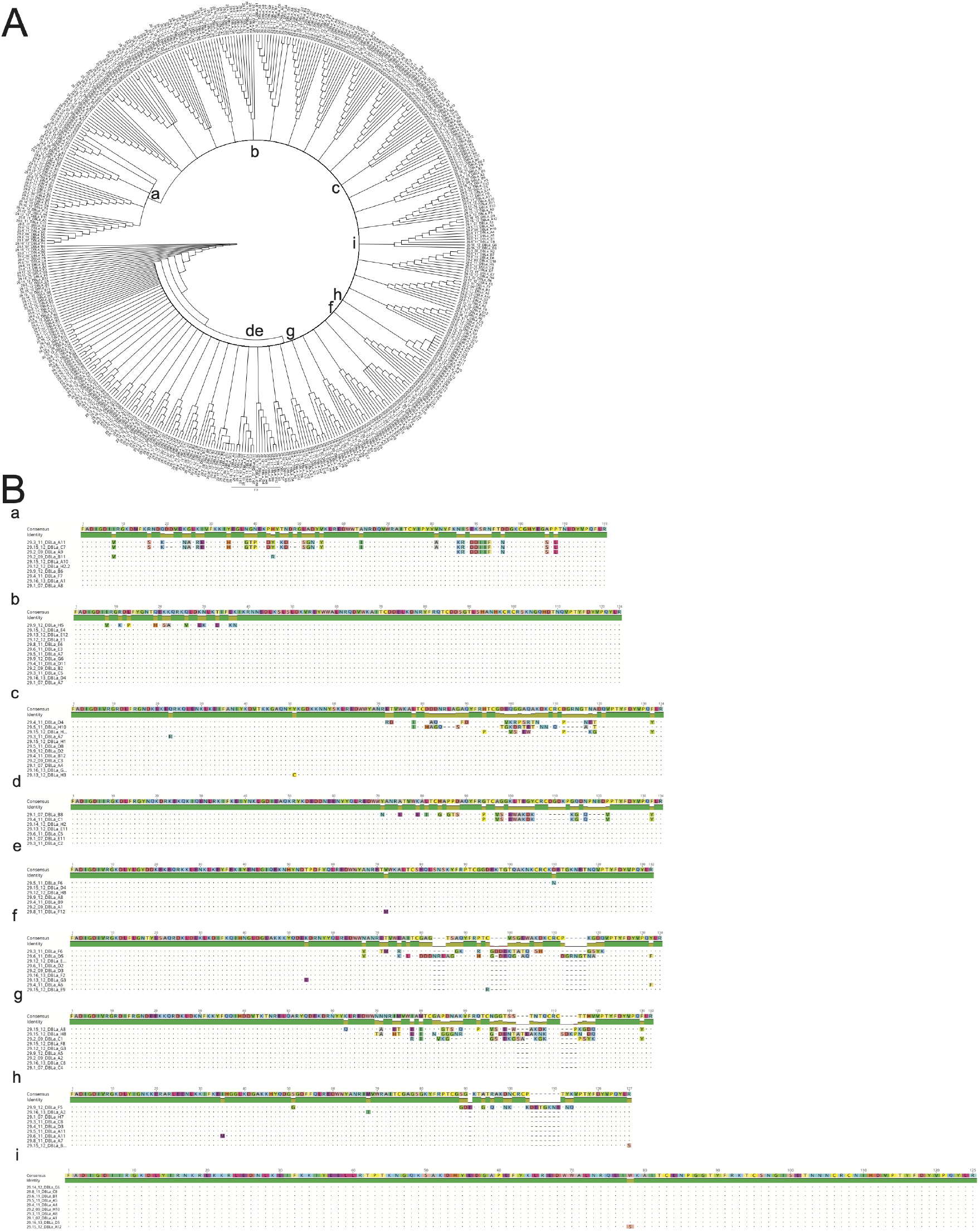
Alignments of Haplotype 29 sequence diversity. A. Neighbor-joining Phylogenetic tree using the Jukes-Cantor method was created using Geneious (version 11.0.2, created by Biomatters) for all DBL1*α* sequences for Haplotype 29. Trees represent a consensus of 1000 bootstrapped replicates, showing nodes present in at least 70% of the replicate trees. Lowercase letters indicate specific branches for which alignments are shown in panel B. B. Sequence alignments of selected branches of the phylogenetic tree are shown. These alignments show different types of diversity for members of the Haplotype 29 cluster, from individual SNPs (as in e, h, i) to blocks of non-homology likely representing var recombination events, evidenced by the presence of a *var* with 100% homology to the other cluster members and an additional var from the same sample with significant diversity in a region of the gene (as in b, c, d, g).

**FIG. S7.**
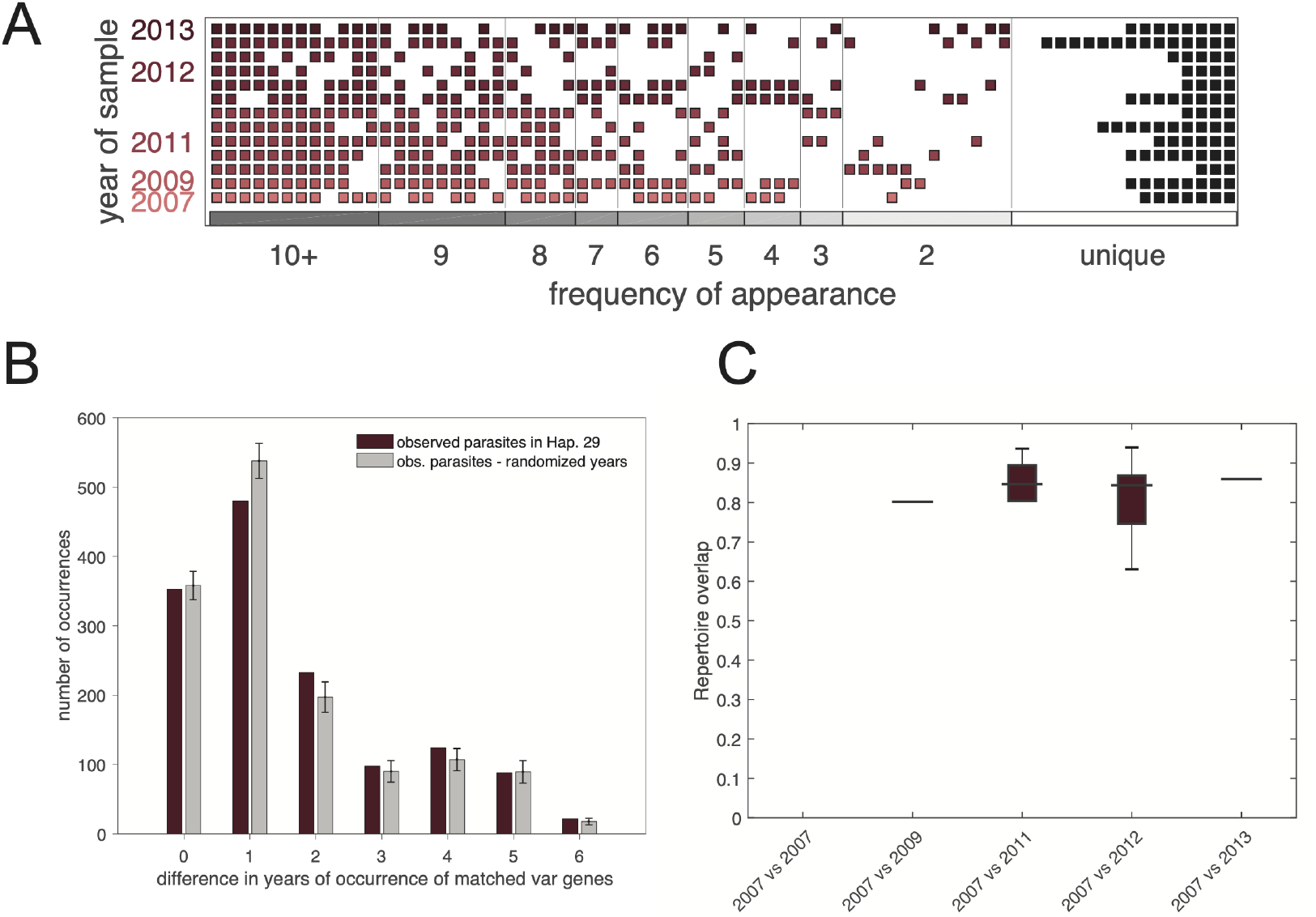
CGS parasites display limited *var* sequence diversification over 7 years. A. Organization of *var* genes in Haplotype 29 over time. Rows represent individual parasite isolates, boxes represent individual *var* genes found in each isolate. Shared *var* genes between years occur in the same vertical column. Black boxes at the far right of the figure represent unique *var* types found in a single isolate. Shared *var* genes are displayed as boxes along the x-axis in order of frequency and are color coded according to isolate. The key at the bottom of the x-axis denotes the frequency (number of isolates sharing a given *var*) in the sampled population. Frequency bin vertical lines have been added for clarity. B. Distribution of differences between the years of observation of matched *var* genes in Haplotype 29 (red) and the same data but with years shuffled (grey; bars indicate means of 1000 independent shuffles; error bars indicate ± one standard deviation). Lack of difference between time-structured and time-randomized distributions indicates lack of time-dependent diversification within Haplotype 29. C. Comparisons of repertoire overlap by year for samples within Hap29. The lack of a linear trend indicates a lack of significant diversification of repertoires over time.

**FIG. S8.**
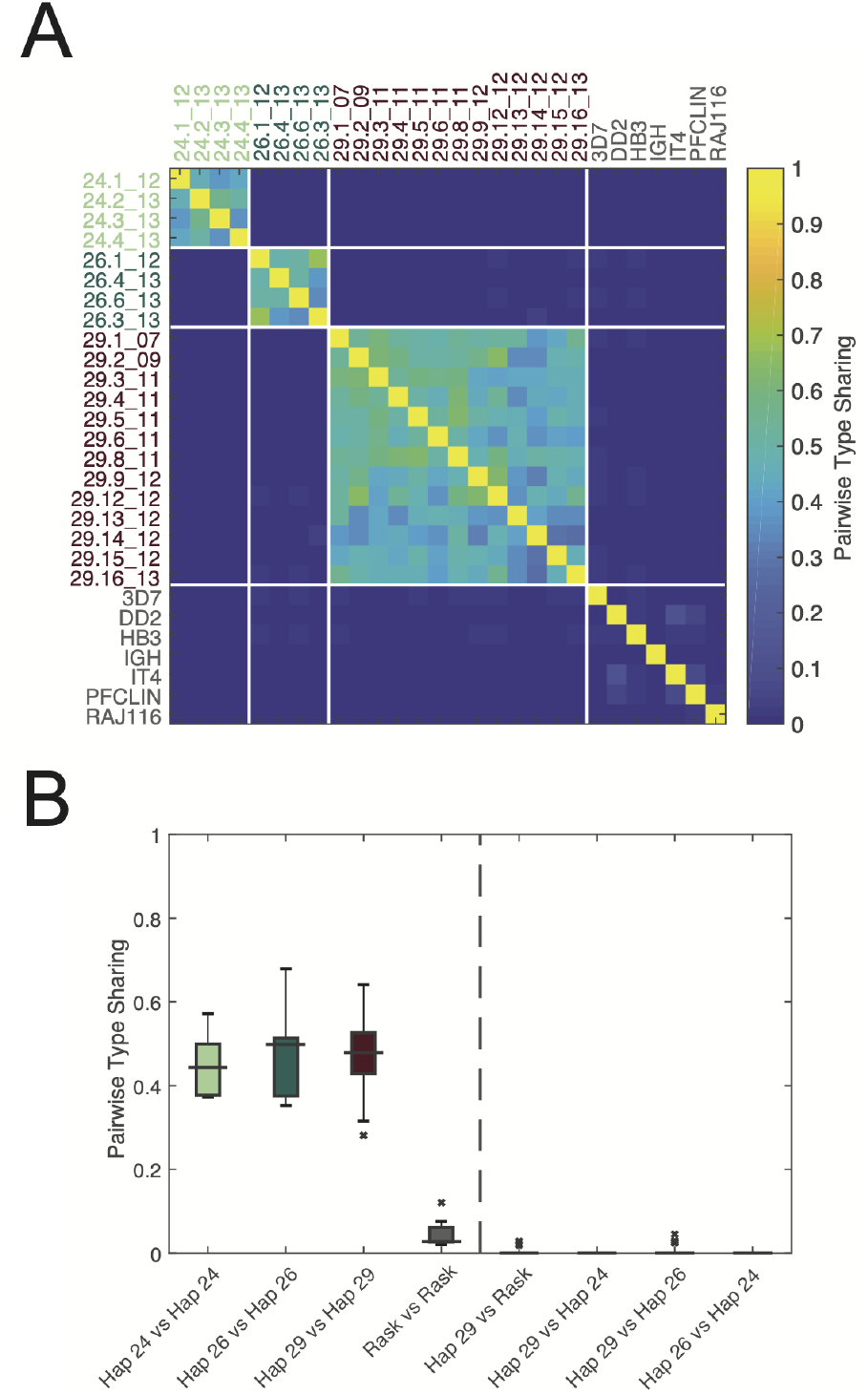
*Var* repertoire overlap of CGS parasites by pairwise type sharing (PTS). Overlap between *var* repertoires among CGS cluster haplotypes 29, 26, and 24, and global sequences (“Rask”), estimated via pairwise type sharing [23], are shown in two ways. A. Heatmap of all-to-all repertoire overlap estimates. B. Boxplots of repertoire overlap distributions within CGS cluster Haplotypes 29, 26, and 24 compared within cluster, between clusters, and to global sequences (“Rask”). Boxes cover interquartile ranges; lines indicate medians; whiskers extend up to 1.5*×*IQR. Identically constructed plots, using the Bayesian repertoire overlap method [19] are shown in Figure 5.

**FIG. S9.**
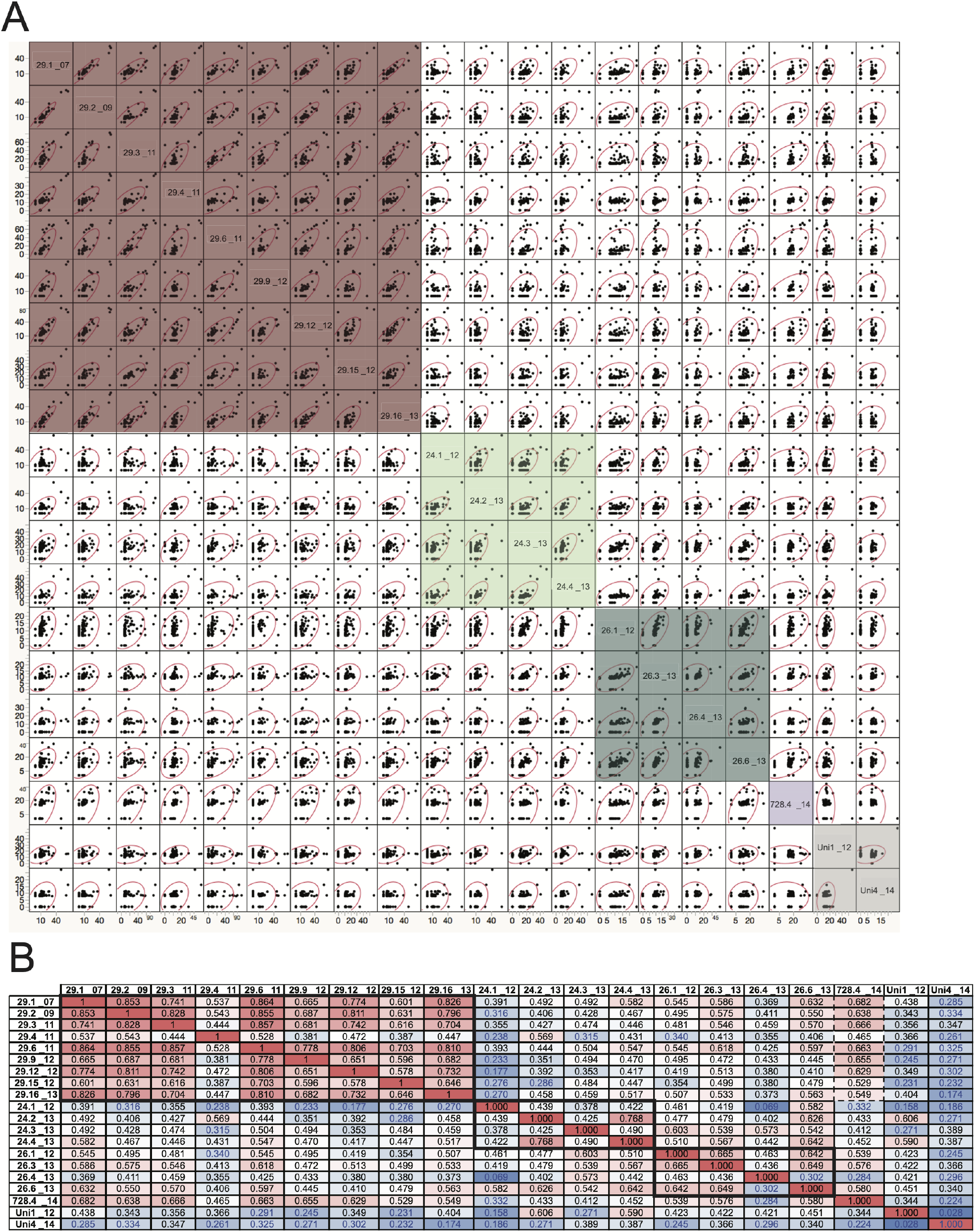
Pairwise comparisons of VSA responses. A. Pairwise comparisons of raw percent IgG recognition from VSA assays for all parasite isolates tested. Colored shaded boxes delineate the clustered haplotypes. B. Correlation coefficients for pairwise comparisons. Individual coefficients are shaded on a scale from low (blue) to high (red). Clustered haplotypes are outlined with a bolded box. Haplotype 728 is outlined with a dotted box to indicate the strong correlation with members of a different Haplotype cluster (Haplotype 29).

**FIG. S10.**
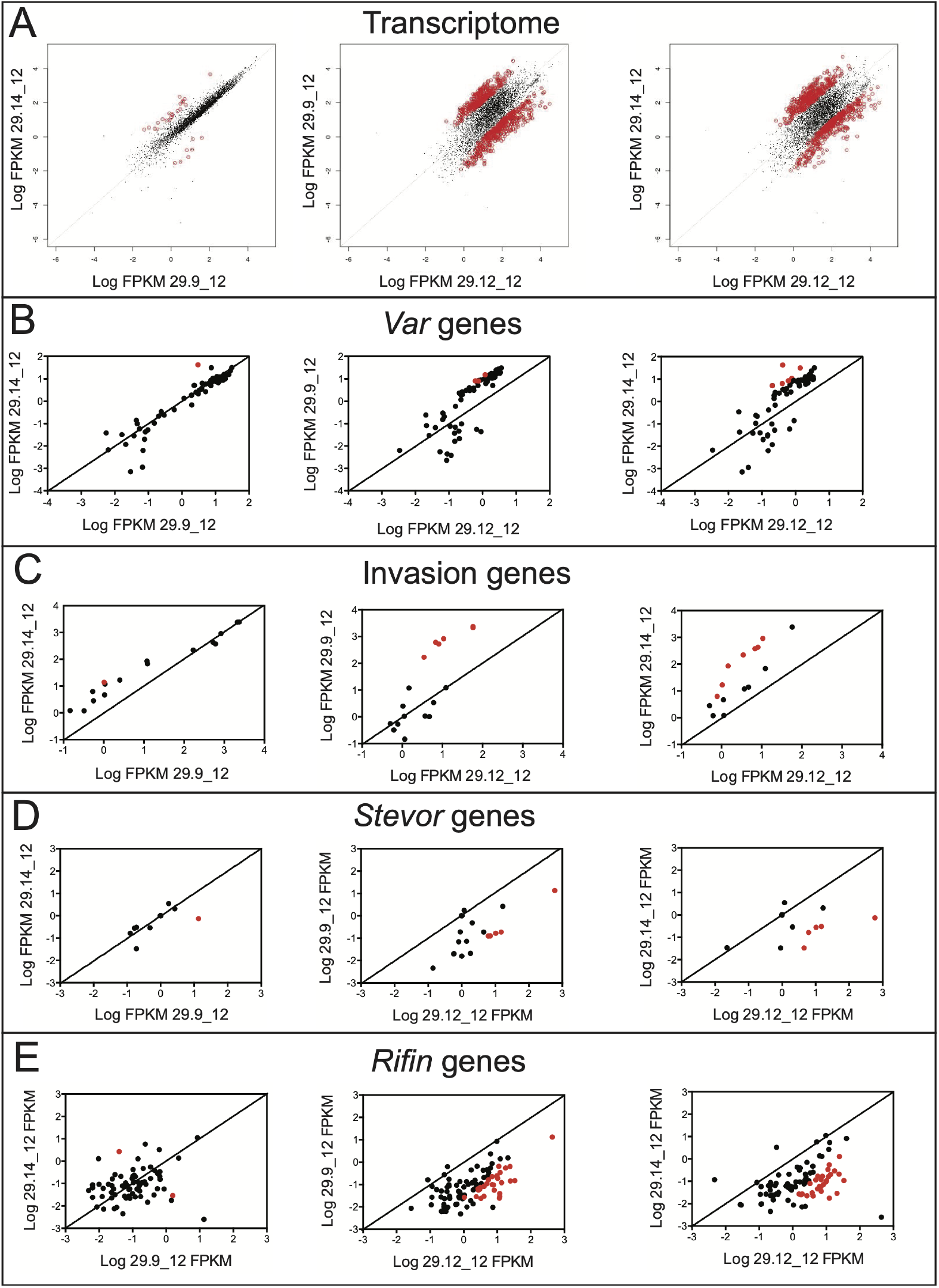
Pairwise comparisons of RNAseq analysis from CGS global and virulence gene transcriptional profiles. log_10_ of FPKM values are shown for all comparisons, with the exception of log_10_ of FPKM values equal to 0. A. Pairwise comparisons of three parasites in Haplotype 29. Globally, the transcriptomes range from being extremely similar (29.9 12 and 29.14 12) to very different (29.9 12 and 29.12 12; 29.12 12 and 29.14 12). Statistically significant differences from paired *t*-tests are indicated as red dots. B. Var gene transcription profiles reveal very similar profiles with few significant differences among 3 isolates within the Haplotype 29 cluster. Statistically significant differences are indicated as red dots. C. Invasion gene transcription profiles reveal very similar profiles with few significant differences among 3 isolates within the Haplotype 29 cluster, with the notable exception of the *Clag* gene family. Statistically significant differences are indicated as red dots. D. *Stevor* gene transcription profiles reveal very similar profiles with few significant differences among 3 isolates within the Haplotype 29 cluster. Statistically significant differences are indicated as red dots. E. *Rifin* gene transcription profiles reveal similar differences among 3 isolates to overall transcriptome comparisons, within the Haplotype 29 cluster. Transcriptional profiles range from being extremely similar (29.9 12 and 29.14 12) to very different (29.9 12 and 29.12 12; 29.12 12 and 29.14 12). Statistically significant differences are indicated as red dots.

**TABLE S1.** Nomenclature for parasite IDs. To simplify the nomenclature for the patient isolates used in the paper, the following convention has been applied: Haplotype.isolate number within the haplotype year.

| Original sample ID | Sample ID code | Haplotype | Year Collected |
| --- | --- | --- | --- |
| Th86.07 | 29.1_07 | 29 | 2007 |
| Th106.09 | 29.2_09 | 29 | 2009 |
| Th103.11 | 29.3_11 | 29 | 2011 |
| Th106.11 | 29.4_11 | 29 | 2011 |
| Th117.11 | 29.5_11 | 29 | 2011 |
| Th123.11 | 29.6_11 | 29 | 2011 |
| Th132.11 | 29.7_11 | 29 | 2011 |
| Th134.11 | 29.8_11 | 29 | 2011 |
| Th114.12 | 29.9_12 | 29 | 2012 |
| Th142.12 | 29.10_12 | 29 | 2012 |
| Th150.12 | 29.11_12 | 29 | 2012 |
| Th162.12 | 29.12_12 | 29 | 2012 |
| Th170.12 | 29.13_12 | 29 | 2012 |
| Th214.12 | 29.14_12 | 29 | 2012 |
| Th230.12 | 29.15_12 | 29 | 2012 |
| Th074.13 | 29.16_13 | 29 | 2013 |
| Th166.12 | 26.1_12 | 26 | 2012 |
| Th203.12 | 26.2_12 | 26 | 2012 |
| Th092.13 | 26.3_13 | 26 | 2013 |
| Th211.13 | 26.4_13 | 26 | 2013 |
| Th245.13 | 26.5_13 | 26 | 2013 |
| Th246.13 | 26.6_13 | 26 | 2013 |
| Th276.13 | 26.7_13 | 26 | 2013 |
| Th068.12 | 24.1_12 | 24 | 2012 |
| Th051.13 | 24.2_13 | 24 | 2013 |
| Th057.13 | 24.3_13 | 24 | 2013 |
| Th061.13 | 24.4_13 | 24 | 2013 |
| Th072.13 | 24.5_13 | 24 | 2013 |
| Th073.13 | 24.6_13 | 24 | 2013 |
| Th095.13 | 24.7_13 | 24 | 2013 |
| Th020.13 | 99.1_13 | 99 | 2013 |
| Th173.13 | 99.2_13 | 99 | 2013 |
| Th178.13 | 99.3_13 | 99 | 2013 |
| Th191.13 | 99.4_13 | 99 | 2013 |
| Th213.13 | 99.5_13 | 99 | 2013 |
| Th102.12 | 81.1_12 | 81 | 2012 |
| Th105.12 | 81.2_12 | 81 | 2012 |
| Th017.13 | 21.1_13 | 21 | 2013 |
| Th036.13 | 21.2_13 | 21 | 2013 |
| Th020.14 | 728.1_14 | 728 | 2014 |
| Th023.14 | 728.2_14 | 728 | 2014 |
| Th024.14 | 728.3_14 | 728 | 2014 |
| Th029.14 | 728.4_14 | 728 | 2014 |
| Th088.12 | Uni1_12 | unique | 2012 |
| Th090.12 | Uni2_12 | unique | 2012 |
| Th208.13 | Uni3_13 | unique | 2013 |
| Th198.14 | Uni4_14 | unique | 2014 |

**TABLE S2.** Sample collection dates and coordinates. Sample collection dates and GPS coordinates are listed for samples in this study. Samples for which ex vivo RNA was available are indicated in bold.

| Strain | Haplotype | Collection Date | Neighborhood | GPS |
| --- | --- | --- | --- | --- |
| 29.1_07 | 29 | 10/11/07 |  |  |
| 29.2_09 | 29 | 10/26/09 |  |  |
| 29.3_11 | 29 | 10/25/11 | Som | 14.780674, -16.937405 |
| 29.4_11 | 29 | 10/27/11 | Medina Fall | 14.806358, -16.909274 |
| 29.5_11 | 29 | 11/2/11 | Escale | 14.797208, -16.929267 |
| 29.6_11 | 29 | 11/4/11 | Lycee Technique | 14.796045, -16.965787 |
| 29.7_11 | 29 | 11/11/11 | Kawsara | 14.808489, -16.915181 |
| 29.8_11 | 29 | 11/14/11 | Keur Cheikh | 14.805962, -16.919905 |
| **29.9_12** | **29** | **9/26/12** | **Diakhao** | **14.798762, -16.938622** |
| **29.10_12** | **29** | **10/5/12** | **Medina Fall** | **14.806358, -16.909274** |
| **29.11_12** | **29** | **10/4/12** | **Mbour 1** | **14.775051, -16.937405** |
| **29.12_12** | **29** | **10/10/12** | **Nguinth** | **14.806814, -16.925882** |
| **29.13_12** | **29** | **10/15/12** | **Penccum Ali Nger** | **14.793105, -16.920345** |
| **29.14_12** | **29** | **11/5/12** | **Medina Fall** | **14.806358, -16.909274** |
| 29.15_12 | 29 | 11/9/12 | Darou Salam | 14.768432, -16.907793 |
| 29.16_13 | 29 | 9/23/13 | Pout | 14.773479, -17.060509 |
| 29.17_14 | 29 | 10/16/14 | Keur Mame Elhadj | 14.801579, -16.924197 |
| **81.1_12** | **81** | **9/21/12** | **Diakhao** | **14.798762, -16.938622** |
| **81.2_12** | **81** | **9/21/12** | **TakhiKaw** | **14.805961, -16.914890** |
| 24.1_12 | 24 | 9/4/12 | Kawsara | 14.808489, -16.915181 |
| **24.2_13** | **24** | **9/16/13** | **Mbal (Pambal)** | **14.967896, -16.895793** |
| **24.3_13** | **24** | **9/17/13** | **Bountou Depot** | **14.794758, -16.918575** |
| **24.4_13** | **24** | **9/18/13** | **Cité Malick Sy** | **14.782373, -16.940837** |
| 24.5_13 | 24 | 9/20/13 | Medina Fall | 14.806358, -16.909274 |
| **24.6_13** | **24** | **9/20/13** | **Dioum (Ndioum)** | **14.776125, -16.943188** |
| **24.7_13** | **24** | **9/25/13** | **Cité Balabé** | **14.794782, -16.913201** |
| 26.1_12 | 26 | 10/11/12 | Thially | 14.804361, -16.936997 |
| **26.2_12** | **26** | **10/30/12** | **Medina Fall** | **14.806358, -16.909274** |
| **26.3_13** | **26** | **9/26/13** | **Hersent** | **14.774989, -16.910197** |
| **26.4_13** | **26** | **10/28/13** | **10éme** | **14.792116, -16.937319** |
| 26.5_13 | 26 | 11/5/13 | Diakhao | 14.798762, -16.938622 |
| 26.6_13 | 26 | 11/5/13 | Diakhao | 14.798762, -16.938622 |
| 26.7_13 | 26 | 11/12/13 | Diakhao | 14.798762, -16.938622 |
| **21.1_13** | **21** | **9/4/13** | **Escale** | **14.797208, -16.929267** |
| **21.2_13** | **21** | **9/11/13** | **Som** | **14.780674, -16.937405** |
| **99.1_13** | **99** | **9/5/13** | **Nguinth** | **14.806814, -16.925882** |
| **99.2_13** | **99** | **10/18/13** | **Bayal (Bayal Khoudia Badiane)** | **14.791653, -16.923446** |
| **99.3_13** | **99** | **10/28/13** | **Diakhao** | **14.798762, -16.938622** |
| **99.4_13** | **99** | **10/23/13** | **Nguinth** | **14.806814, -16.925882** |
| **99.5_13** | **99** | **10/28/13** | **Nassrou** | **14.766740, -16.902761** |
| **728.1_14** | **728** | **9/25/14** | **Diakhao** | **14.798762, -16.938622** |
| **728.2_14** | **728** | **9/25/14** | **Nguinth** | **14.806814, -16.925882** |
| **728.3_14** | **728** | **9/25/14** | **Nguinth** | **14.806814, -16.925882** |
| **728.4_14** | **728** | **9/29/14** | **Diakhao** | **14.798762, -16.938622** |
| **Uni1_12** | **Unique** | **9/17/12** | **Diakhao** | **14.798762, -16.938622** |
| **Uni2_12** | **Unique** | **9/17/12** | **10éme** | **14.792116, -16.937319** |
| **Uni3_13** | **Unique** | **10/28/13** | **Cité Balabé** | **14.794782, -16.913201** |
| **Uni4_14** | **Unique** | **11/18/14** | **Diakhao** | **14.798762, -16.938622** |

**TABLE S3.**
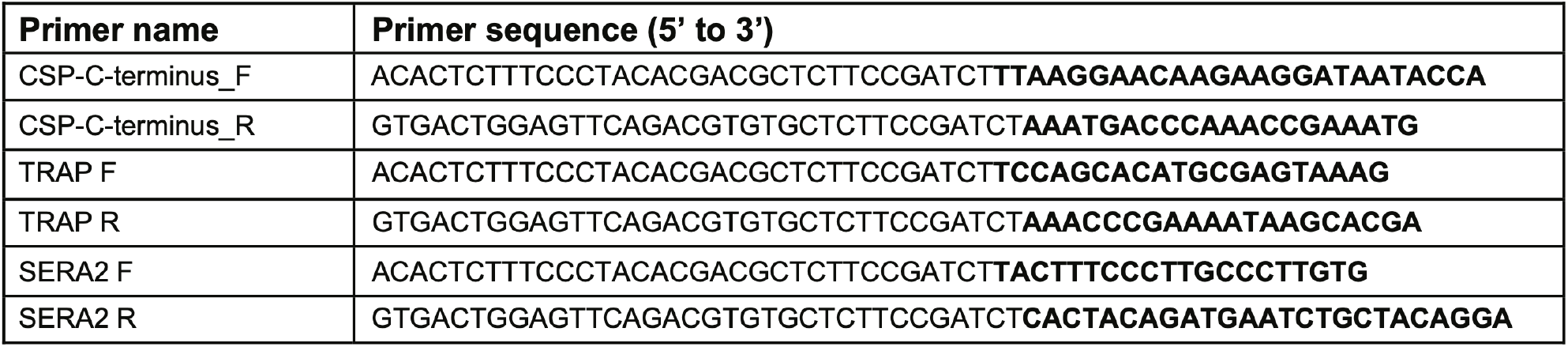
Primer sequences for CSP, TRAP, and SERA2. Primer sequences used for PCR amplification and sequencing of CSP C-terminus, TRAP, and SERA2 are shown in the 5’ to 3’ orientation with the *Plasmodium* portion of the primer sequence noted in bold.

